# A computational model of *Pseudomonas syringae* metabolism unveils the role of branched-chain amino acids in virulence expression at the early stages of Arabidopsis colonization

**DOI:** 10.1101/2022.12.16.520825

**Authors:** Philip J. Tubergen, Greg Medlock, Anni Moore, Xiaomu Zhang, Jason A. Papin, Cristian H. Danna

## Abstract

Leaf mesophyll-colonizing bacterial pathogens infect their plant hosts by adjusting their metabolism to the leaf mesophyll environment. Soon after the inoculation of naïve, susceptible plants, the model bacterial pathogen *Pseudomonas syringae* pv. *tomato* DC3000 (*Pst*DC3000) expresses virulence factors that suppress plant immunity, a requirement to produce robust infections. However, if plant immunity was elicited with Microbe-Associated Molecular-Patterns (MAMPs) prior to bacterial inoculation, *Pst*DC3000 slows down virulence expression and only produces symptomless mild infections. To understand how bacterial metabolism adapts to these two contrasting conditions, we created iPst19, an *in silico* ensemble of genome-scale metabolic reconstructions. Constraining the *in silico* growth of iPst19 with *in planta Pst*DC3000 gene expression data revealed that sugar catabolism is highly active in bacteria that have been inoculated in mock-treated plants. In contrast, branched-chain amino acids (BCAAs) catabolism is highly active in bacteria that have been inoculated in MAMP-pretreated plants. Bacterial growth and gene expression analysis showed that BCAAs suppress virulence gene expression without affecting bacterial growth *in vitro. In planta*, however, BCAAs suppress the expression of virulence genes at the early stages of the infection and significantly impair leaf colonization of the host plant *Arabidopsis thaliana*. While the overexpression of the conserved bacterial leucine-responsive transcriptional regulator *Lrp* induced the expression of virulence genes, its downregulation had the opposite effect, suggesting that BCAA-free Lrp induces virulence while BCAA-*Lrp* does not. Overall, our data provide mechanistic connections to understand how plant immunity impacts *Pst*DC3000 metabolism and virulence, furthering our understanding of bacterial pathogenesis and plant disease.

## INTRODUCTION

*Pseudomonas syringae* is a pathogenic bacterium that can infect a wide range of plant species, often in a species-specific interaction (Xin and He, 2013). Based on its host range, more than 60 different pathovars have been identified (Gomila et al., 2017). In addition to causing disease in tomatoes, *Pseudomonas syringae* pv. *tomato* DC3000 (*Pst*DC3000) can infect the model plant *Arabidopsis thaliana* (Whalen et al., 1991). The *Pst*DC3000-Arabidopsis pathosystem is genetically tractable; thus, it is a preferred model that has facilitated the discovery of molecular mechanisms underlying plant defense and bacterial disease that have been translated to distant plant species, including crops (Xin and He, 2013).

*Pst*DC3000 is an endophytic pathogen that multiplies in the intercellular spaces of the leaf mesophyll, also called leaf apoplast (LA). The LA is partially filled with an aqueous solution rich in proteins, sugar polymers, free hexoses, free amino acids, and other plant-synthesized metabolites that can support *Pst*DC3000 growth (Rico and Preston, 2008). The concerted activity of membrane transporters and LA resident enzymes regulate the concentration of each metabolite in basal conditions and in response to stress (Lopez-Millan et al., 2000; Sattelmacher, 2001). Once inside the LA, *Pst*DC3000 subverts early inducible plant defense responses that would otherwise restrict its growth and prevent the onset of disease (Crabill et al., 2010; Lovelace et al., 2018; Nobori et al., 2018). *Pst*DC3000 uses two main virulence factors to suppress plant immunity and achieve colonization: the Type-3 Secretion System (T3SS) that injects effector proteins into host cells, regulated by the transcriptional regulator *hrpL*, and the phytotoxin coronatine, which requires the enzyme coronafacate ligase (*cfl*) for its synthesis (Rangaswamy et al., 1997; Guo et al., 2009; Crabill et al., 2010). Modulation of these virulence factors has been previously described as dependent on nutritional and environmental cues. Plants perceive invading *Pst*DC3000 via the recognition of conserved molecules generically known as Microbe-Associated Molecular Patterns (MAMPs). This recognition is mediated by Pattern-Recognition Receptors (PRRs) at the plasma membrane. PRRs can be activated with purified synthetic MAMPs prior to bacterial infection. Such activation elicits changes in the composition of the LA (Zhang et al., 2022b; Zhang et al., 2022a), which in turn, suppress the expression of *Pst*DC3000 T3SS genes, thus lowering *Pst*DC3000 virulence and infectivity (Anderson et al., 2014; Yamada et al., 2016; Zhang et al., 2022b). Altogether, these studies demonstrate that plant defense responses drive changes in the composition of metabolites that contribute to inhibiting *Pst*DC3000 growth and preventing the onset of plant disease. Notwithstanding the relevance of these discoveries, they likely represent only a glimpse of the contribution of LA metabolites to plant defense.

Amino acids are emerging as important regulators of *Pst*DC3000 virulence within the LA. Broadly, the nutritional state of the environment plays a key role in the virulence response of pathogenic bacteria. This seems especially true for *Pst*DC3000, as amino acid supplementation into the LA during infection with *Pst*DC3000 decreases long-term leaf colonization in an AA-specific manner. *Pst*DC3000 shows decreased colonization of the leaf after 72 hours when co-infiltrated with either glutamine, serine, or valine (Zhang et al., 2022b). Branched-chain amino acids (BCAAs), including valine, have been shown to regulate virulence in gram-negative bacteria by modulating the activity of the global transcriptional regulator *leucine-responsive regulatory protein* (*Lrp*) (Kaiser and Heinrichs, 2018). *Lrp* activity responds to the bacterial nutritional state and regulates both virulence and the transition into the stationary phase. In low BCAA concentrations, *Lrp* induces the expression of the acetolactate synthase gene encoding an enzyme that uses pyruvate as a substrate for the synthesis of BCAAs (Subashchandrabose et al., 2009). In addition, *Lrp* regulates the synthesis of the pili, among other virulence factors, the transition into the stationary phase, and amino acid metabolism in *E. coli* and *Salmonella sp*. (Cho et al., 2008)(Baek et al., 2009). The full spectrum of regulations mediated by *Lrp* is not fully understood, yet it is clear that amino acids, particularly BCAAs, control which genes are regulated by *Lrp* (Cho et al., 2008; Kroner et al., 2019).

The complex composition of the LA and the dynamic changes in the concentration metabolites that take place during the course of infections (Solomon and Oliver, 2002; Ward et al., 2010; O’Leary et al., 2016) have hindered efforts to define which plant metabolites, and at which concentrations, would have a positive or a negative impact on *Pst*DC3000 growth *in planta*. To capture this complexity, recently published studies have used bacterial gene expression profiling to understand how the LA affects *Pst*DC3000 growth and virulence (Lovelace et al., 2018; Nobori et al., 2018). These studies hypothesized that MAMP-elicited immunity could suppress the expression of bacterial virulence genes through metabolite deprivation, as previously reported by Anderson and colleagues (Anderson et al., 2014). In addition, Nobori and colleagues (Nobori et al., 2018) found that plant immunity also suppresses the expression of genes encoding ribosomal proteins, suggesting that MAMP-treated plants may broadly impact *Pst*DC3000 protein synthesis as well. Besides agreeing on *Pst*DC3000 virulence suppression, these two studies reached dissimilar conclusions with regard to the plant metabolites that may restrict *Pst*DC3000 growth *in planta*. This apparent discrepancy may stem from experimental differences across these studies. However, it is also possible that *Pst*DC3000 transcriptomics analyses alone are insufficient to capture the complexity of the metabolic changes that define *Pst*DC3000 growth *in planta*. To overcome these potential limitations, we have generated an ensemble of **ge**nome-scale **n**etwork **re**constructions (GENREs) and hypothesize that metabolic modeling could capture previously undescribed metabolic shifts that multi-omics alone cannot contextualize.

GENREs, and their corresponding modeling counterpart, called **ge**nome-based **m**odels (GEMs), have emerged as a powerful tool for predicting metabolic phenotypes and gene essentiality (Blazier and Papin, 2012). GENREs are originally built from genome annotations and are curated with various forms of evidence, including *in vitro* metabolic demands, protein homology, and literature research (Thiele and Palsson, 2010). Within a GEM, transport reactions facilitate the movement of metabolites from one compartment (i.e., the outside environment) to another compartment (i.e., inside the cell). Gene expression data can be overlaid on top of a GEM framework to make integrative predictions of metabolic activity. Multiple algorithms for integrating gene expression data with GEMs have been developed, each of which makes assumptions about the relationship between gene expression and reaction activity (Richelle et al., 2019).

Here we have developed a metabolic model of *Pst*DC3000, iPst19, using the well-annotated genome of *Pseudomonas syringae* pv. *tomato* DC3000 (Buell et al., 2003). Through literature mining, sequence homology comparisons, and ensemble gap-filling (Medlock and Papin, 2020), we have iteratively curated the draft reconstruction to be more akin to *Pst*DC3000 biology. Leveraging the unbiased systems-level view, iPst19 made new and insightful predictions that were confirmed experimentally *in vitro* and *in planta*.

## RESULTS

### iPst19, an ensemble of genome-scale metabolic models

A draft reconstruction of the *Pst*DC3000 GENRE was generated using ModelSEED (Henry et al., 2010) and genome data from Buell and colleagues (Buell et al., 2003). We modified the draft reconstruction using homologous comparison with the published Pseudomonas GEMs iPae1146, and iPau1129, representative of *Pseudomonas aeruginosa* strains PAO1 and PA14, respectively. The draft was further curated through ensemble gap-filling (**Figure 1A**). To gap-fill the GEM, we used an approach called **A**utomated **M**etabolic **M**odel **E**nsemble-**D**riven **E**limination of **U**ncertainty with **S**tatistical learning (AMMEDEUS) (Medlock and Papin, 2020). Briefly, AMMEDEUS incorporates traditional gap-filling of metabolic models based on *in vitro* growth data on single carbon sources (SCS), identifying reactions that collectively carry the minimal amount of flux necessary to produce biomass *in silico* in each condition. To create a carbon utilization list for AMMEDEUS, we performed SCS growth phenotyping using Biolog phenotype microarrays PM1 and PM2a plates (Hayward, CA). We grew *Pst*DC3000 in each of 190 SCS in quadruplicate and recorded the optical density at 600nm (OD600) at 0, 12, 24, 36, 48, and 60 hours (**Figure 1B**). To only include high-confidence growth conditions for gap-filling, we only considered conditions that resulted in a max OD600 greater than 0.1 after subtraction of the 0-hour baseline. Following AMMEDEUS, gap-filled reactions that presented the most uncertainty for producing flux through the biomass function in a complete media were assessed for literature support in *Pst*DC3000 and other Pseudomonads. Reactions with literature evidence were added to the reconstruction. This iterative process was completed three times to form the current 100-member ensemble iPst19 (Figure 1D). Several amino acids and sugars produced significant growth over 60 hours of incubation **(Figure 1B**). Of special interest were plant metabolites previously shown to have an impact on *Pst*DC3000 infections (Rico and Preston, 2008; Park et al., 2010; Anderson et al., 2014; Zhang et al., 2022b). GABA, a highly abundant amino acid in the LA of tomato plants, produced robust growth, as did L-glutamine, sucrose, and D-glucose (**Figure 1B**). We then assessed the *in silico* biomass production on SCS to preliminarily validate the ensemble. Of the highlighted substrates in **Figure 1B**, all *in silico* simulations were able to predict the *in vitro* growth outcome (**Figure 1C**). The resulting model comprises 1439 unique reactions and 1224 unique metabolites, while each individual ensemble contains 1330 +/- 7 reactions, 1215 +/- 2 metabolites, and 889 genes. iPst19 is similar in size to the well-curated *Pseudomonas aeruginosa* GEMs and is smaller than the gold-standard *E. coli* GEM, likely due to the depth of experimental support available for *E. coli* (**Figure 1D**). To estimate the success of the curation process, we assessed the composition of the annotated genes included in iPst19 for metabolic relevance. Some genes have multiple annotated ontologies, resulting in a higher number of total ontologies than genes, with 437 genes annotated with ontologies related to the metabolism of amino acids, followed by 335 genes related to secondary metabolism and 224 antibiotic metabolism-related genes (**Figure 1E**).

**Figure 1.**
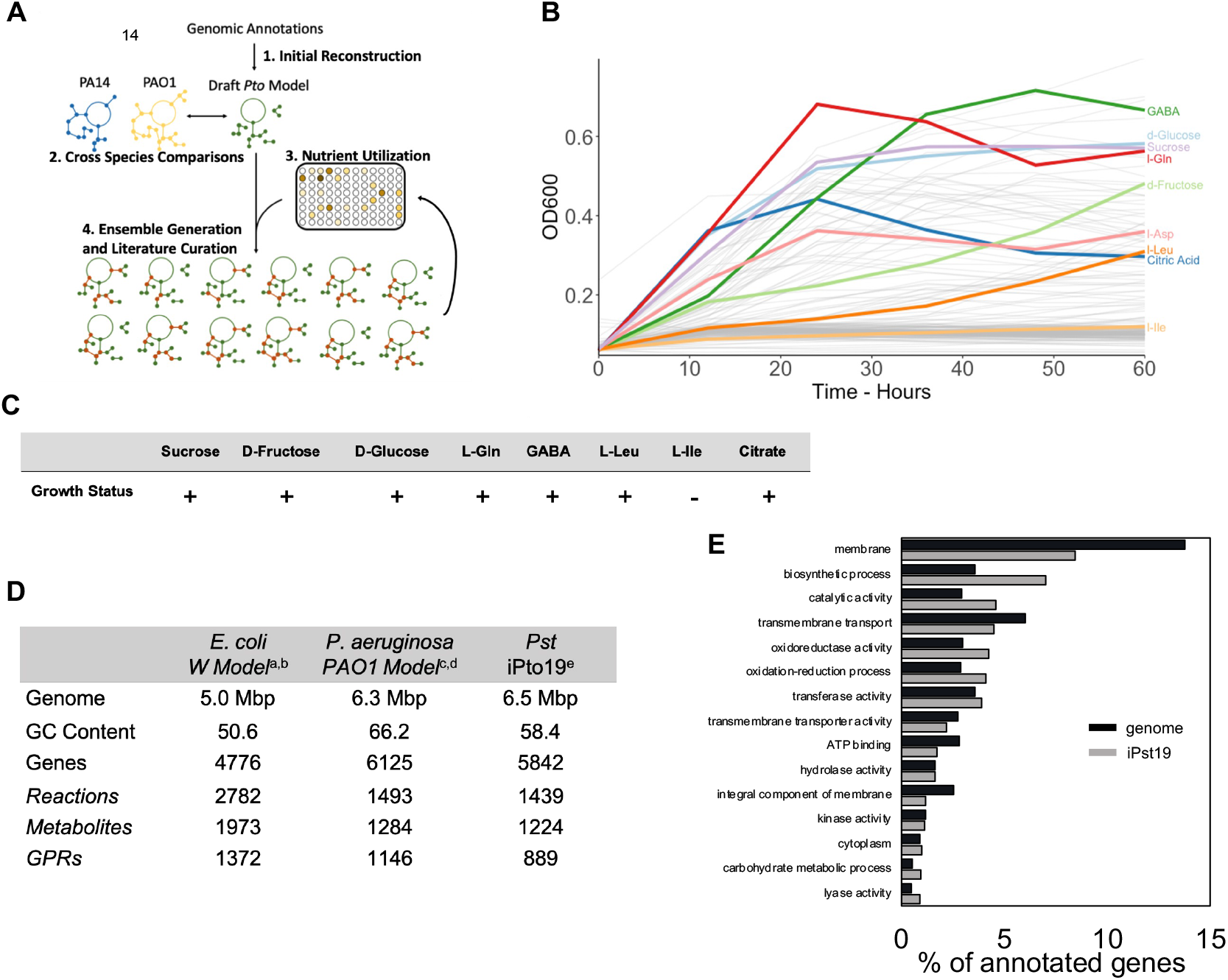
Construction and validation of iPst19. **(A)** Generation of iPst19, including the initial reconstruction from gene annotations, comparison to other *Pseudomonas* models, integration of SCS growth data, and iterations of gap-filling and manual curation. **(B)** OD600 for Pst grown in 190 SCS media over 60 hours with constant agitation (grey lines). Shown in color are SCS metabolites found in the leaf apoplast of Arabidopsis plants. **(C)** Output of SCS *in silico* growth simulations using iPst19. **(D)** Comparison of the size of well-curated models and the corresponding genomes. Genome information for Escherichia coli (^a^) Pseudomonas aeruginosa (^c^) and Pseudomonas syringae (^e^) were acquired from publicly available genome assemblies. (a) (Archer et al., 2011) (c) NCBI Tax ID 187; (e) (Buell et al., 2003). (b) The *E. coli* W model was obtained from BiGG (King et al., 2016) (Monk et al., 2013). (d) The *P. aeruginosa* model was created by Oberhardt et al. (Oberhardt et al., 2008). **(E)** Gene orthologies in iPto19 show biosynthetic processes and catalytic activities. GO terms for genes within iPto19 show enrichment of metabolically-focused orthologies when compared to the whole PstDC3000 genome.

### Gene expression-constrained metabolic flux highlights the role of leucine catabolism under growth-restrictive conditions imposed by plant immunity

Integration of transcriptomic profiles into a GEM can contextualize complex relationships between genes and corresponding metabolic pathways (Blazier and Papin, 2012). To study the metabolic changes that *Pst*DC3000 experiences in the LA, we use the GIMME integration algorithm (Becker and Palsson, 2008) to constrain iPst19 metabolic flux with a previously published *in planta* gene expression dataset of *Pst*DC3000 obtained 5h post-inoculation into leaves of Arabidopsis plants (Nobori et al., 2018). Of the 889 genes included in iPst19, 250 were significantly up or down-regulated in *Pst*DC3000 inoculated in flg22-treated plants compared to mock-treated plants (**Dataset S1**). With the gene expression-constrained metabolic flux, we identified reactions that carried the most disparate flux between the two contrasting conditions analyzed: mock-treated plants, where *Pst*DC3000 grows aggressively, versus flg22-pretreated plants, where *Pst*DC3000 grows modestly. Among the twenty reactions with the most contrasting fluxes (**Figure 2**), the catabolism of leucine (reaction no. 9 and no. 17) showed increased flux in flg22-elicited plants where bacterial growth is compromised. In *Pst*DC3000, both reactions are associated with *LiuA*, a unique enzyme with isovaleryl-CoA dehydrogenase activity encoded by PSPTO_2739 (**Table S1; Dataset S2**). In contrast to BCAAs degradation, sugar-related catabolic reactions were disproportionately active in *Pst*DC3000 inoculated in mock-treated plants. To understand the biological function of these metabolic signatures, we sought to run an *in-silico* genetic screen to identify genes essential to the utilization of glucose and leucine.

**Figure 2.**
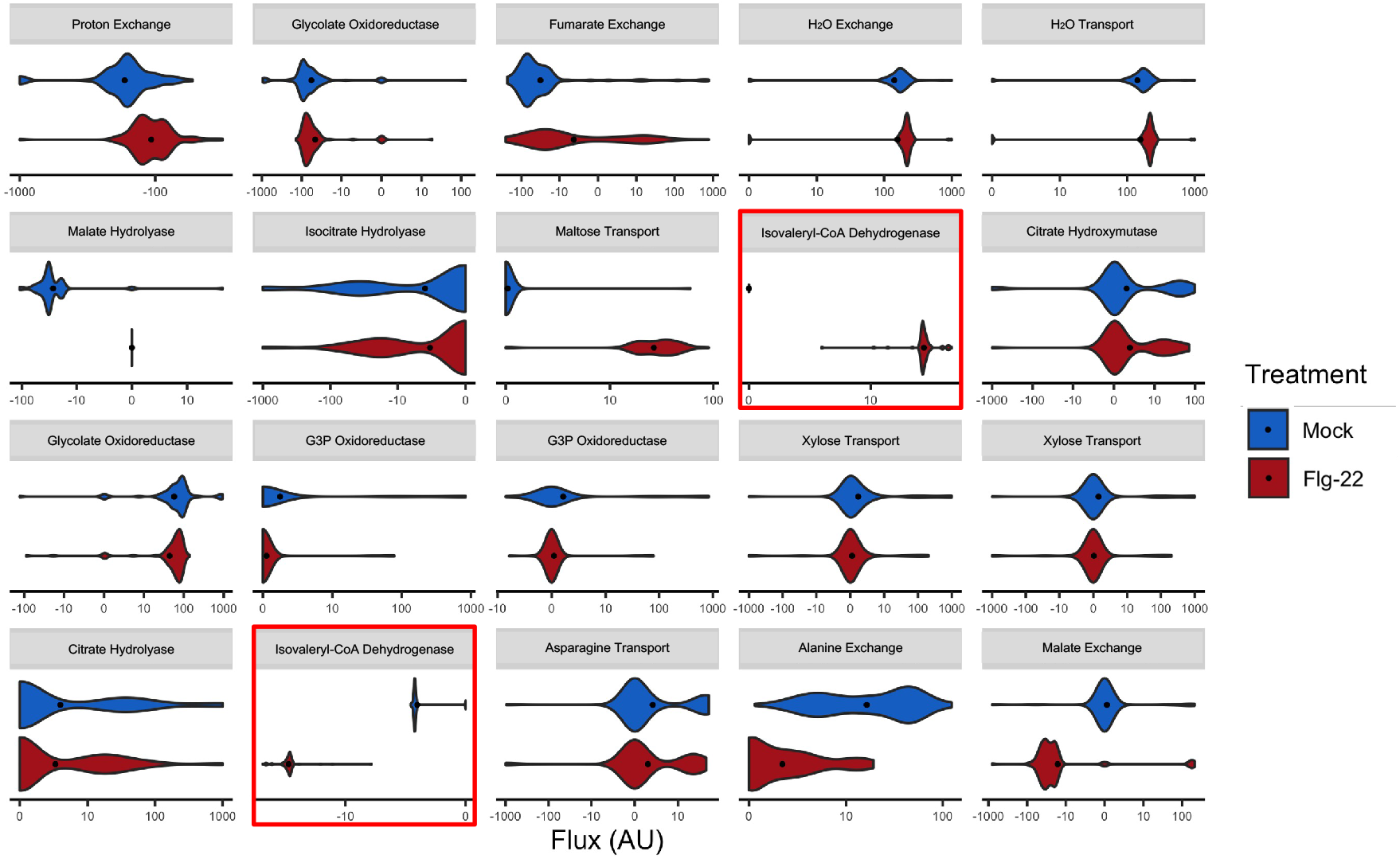
*Pst*DC3000 gene expression-constrained iPst19 metabolic flux. RNAseq datasets (Nobori et al., 2019) were integrated into iPst19 to constrain fluxes of reactions using the GIMME integration algorithm. Each of the 20 reactions with the most discordant fluxes between constrained conditions is presented for comparison. The black dot represents the mean flux of all 100 ensemble members for the reaction. Constraints generated from PstDC3000 gene expression in mock-treated or flg22-treated plants is presented in blue and red, respectively. Reactions number 9 and 17, whose fluxes are constrained by PSPTO_2739, are highlighted with red squares.

### iPst19 predicts the essentiality of the *liu* operon to metabolize leucine

Within a metabolic model or ensemble, gene essentiality varies depending on the substrates available for growth simulation. Due to the gap-filling process used for optimizing iPst19, each of the 100-member ensembles has a unique architecture. Therefore, we simulated single gene knockouts across the whole 100-member ensemble and scored essentiality as the proportion of members of the ensemble that predict a given gene as essential depending on the carbon source available. First, and informed by data obtained from constraining iPts19 growth with *Pst*DC3000 transcriptional data, we tested gene essentiality in three different media: a complete media, a glucose minimal media, and a leucine minimal media. Glucose and leucine were tested to serve two purposes: glucose alone produces robust growth experimentally yet lacks a nitrogen moiety itself, while leucine produces more modest growth experimentally, but provides a nitrogen moiety, thus potentially relieving reliance on ammonium as the sole source of nitrogen in the glucose-only medium. From a total of 889 genes, the analysis identified 136 predicted essential genes necessary for biomass production in a rich (complete) medium (**Figure 3A**). In addition, we identified 28 substrate-specific predicted essential genes (**Figure 3B**). While twenty-three genes were common to both glucose and leucine SCS media, five were differentially essential. The leucine minimal medium rendered additional genes predicted to be essential: PSPTO_2736, PSPTO_2738, and PSPTO_2739, each of which encode enzymes of the leucine catabolic pathway (**Figure S1**). BCAAs degradation requires α-ketoacid dehydrogenase enzymatic complexes composed of three subunits (E1, E2, and E3), each with different reaction specificities. In *Pseudomonas putida*, the leucine catabolism begins with the conversion of leucine into 4-methyl 2-oxopentanoate, a reaction performed by a BCAA-aminotransferase. In *Pst*DC3000, this enzyme is encoded by PSPTO_1332. The next step in leucine degradation requires branched-chain keto acid dehydrogenase (BCKDC) enzymes. While in *P. putida*, these enzymes are encoded by the *bkd* operon (Marshall and Sokatch, 1972), the *Pst*DC3000 genome lacks an operon encoding BCKDCs. BCKDC enzymes, together with those of the 2-oxoglutarate dehydrogenase complex (ODHC) and the pyruvate dehydrogenase complexes (PDC), belong to the α-ketoacid dehydrogenase protein family and share high percentages of identity with one another (Burns et al., 1988; Hester et al., 1995). Notwithstanding the lack of a *bkd* operon, *Pst*DC3000 can use leucine as an SCS (Figure 1B). Hence, we hypothesize that ODHC or PDC enzymes could take 4-methyl 2-oxopentanoate to produce isovaleryl-CoA, the substrate of *LiuA* (**Figure S1**). To identify enzymes with BCKDC-like activity in *Pst*DC3000, we use the amino acidic sequences of enzymes encoded by the *P. putida bkd* operon as a query in BLASTp (https://blast.ncbi.nlm.nih.gov/). The search yielded one ODHC-E1 component encoded by PSPTO_2199, one ODHC-E2 component encoded by either PSPTO_2200 or PSPTO_5006, and one ODHC-E3 component encoded by PSPTO_2201 (**Table S2**), suggesting that enzymes of the ODHC complex could be part of the leucine degradation pathway and serve function equivalent to those of the BCKDC complex in *P. putida*.

**Figure 3.**
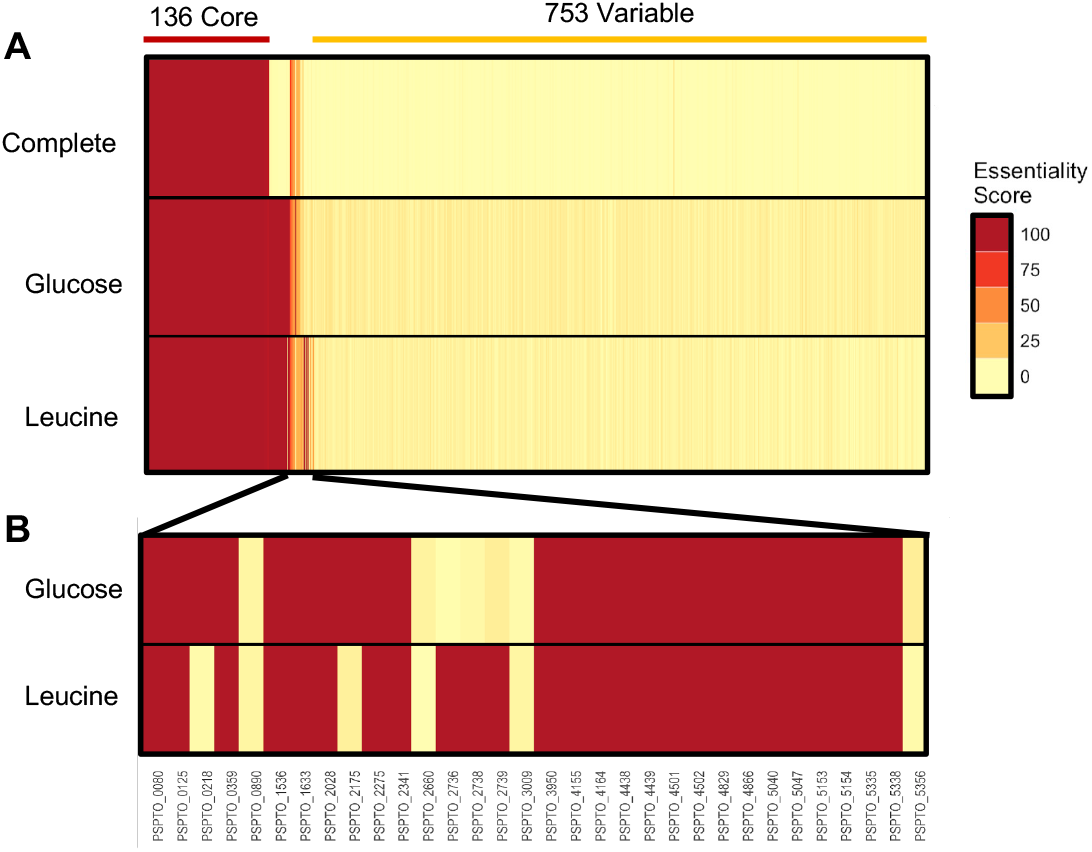
iPst19 predicts genes of the *liu* operon as essential for Leu catabolism. Each of the 889 genes present in iPst19 was removed from a total of 100 ensembles one at a time, and final biomass production was assessed. (**A)** Predicted essentiality in each medium is given out of 100 ensemble members. An essentiality score of 100 indicates essentiality in all 100 ensemble members, while a score of 0 indicates non-essentiality in all 100 ensembles. iPst19 detected 136 shared predicted essential genes across all tested conditions. (**B)** Twenty-three predicted glucose- and leucine-specific essential genes are shown, five of which have varying degrees of essentiality depending on the carbon source used. Three genes of the *liu* operon (PSPTO_2736, PSPTO_2738, and PSPTO_2739) are essential to use leucine as the sole carbon source.

The glucose minimal medium rendered two genes differentially essential from the leucine minimal medium: PSPTO_0218, an ammonium transporter encoding gene, and PSPTO_2175, a gene coding for the iso-propyl-malate dehydrogenase activity that catalyzes one of the first steps in BCAA biosynthesis. Growth assessment in SCS media provides valuable information on metabolite utilization but does not mimic the complex metabolite composition of the LA. To analyze gene essentiality in a two-substrate environment, we simulated growth and generated a gene essentiality profile on media containing D-glucose and L-leucine in varying concentrations. The total availability of mixed substrates in the media was maintained at 10 mM/gDW. In a mixed substrate with 99% glucose and 1% L-leucine, gene essentiality matches predicted gene essentiality in D-glucose alone. Similarly, with 1% D-glucose and 99% L-leucine, the essentiality profile resembles that of L-leucine alone. As expected, the essentiality profile of 50% D-glucose and 50% L-leucine, as well as those with 10% of a second substrate, included fewer genes than either single substrate (**Figure 4A**). The essentiality of genes was not a binary output immediately alleviated by the introduction of the second metabolite. Instead, there is a predicted threshold at which leucine alleviates the need for glucose-only essential genes and *vice versa* (**Figure 4B**). Since these genes are predicted to be essential only when leucine content reaches 99% of the carbon source in the medium, our results suggest that the concentration of leucine alone would not explain the expression patterns of *liuA* and *liuD in planta* (**Dataset S1**). In addition, the data suggest that BCAA degradation may not contribute significant amounts of acetyl-CoA to the tricarboxylic acid (TCA) cycle. Instead of supporting bacterial growth, *liuA* and *liuD* may play a regulatory role during the infection process by modulating the levels of leucine and other BCAA that serve as signals to adjust bacterial growth to changing environmental conditions in the LA.

**Figure 4.**
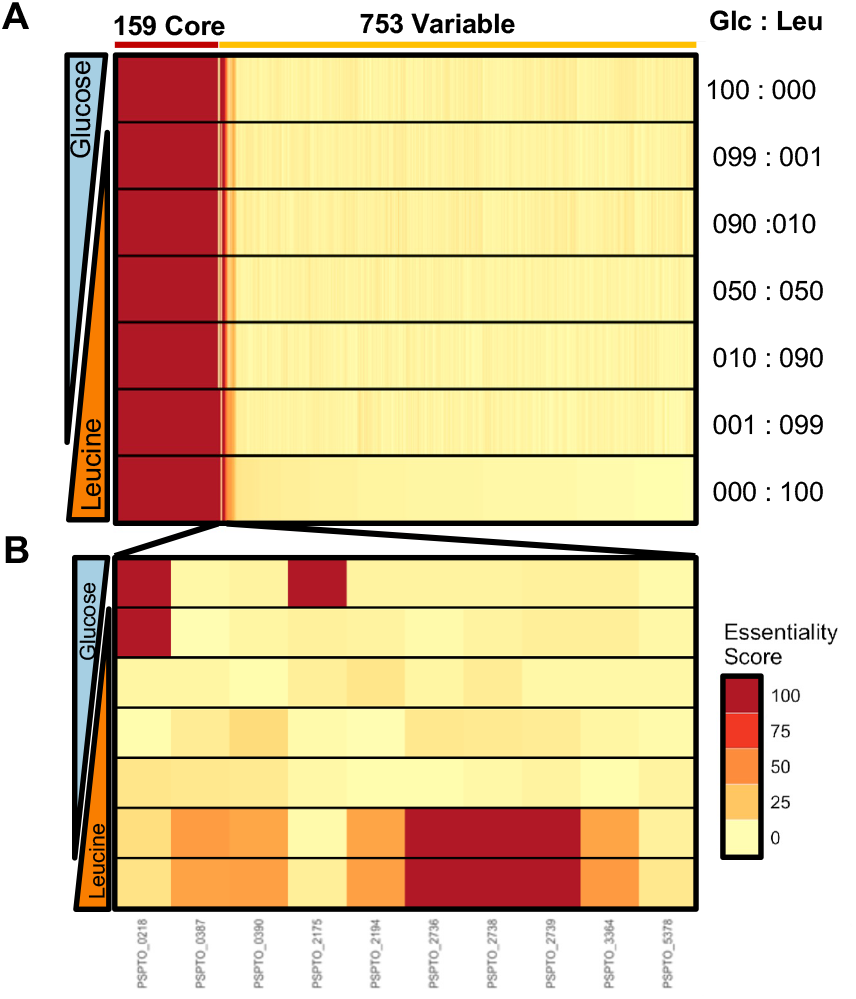
Predicted gene essentiality changes with mixed carbon sources. iPst19 gene essentiality was simulated in conditions where carbon sources were restricted to an uptake rate of 10 mM/g dry weight. The carbon composition of the Glc and Leu simulated media is indicated to the right as a percentage. (**A)** All genes present in iPst19 were simulated for essentiality. **(B**) The most dissimilar essential genes in D-glucose and L-leucine simulated medium.

### BCAAs contribute to the regulation of virulence expression *in vitro* and *in planta*

*P. aeruginosa* controls leucine levels by a feedback loop where high leucine levels suppress leucine biosynthesis and induce leucine degradation, while low leucine levels have the opposite effect (Rabin et al., 1968; Aguilar et al., 2006; Cattoir et al., 2012). The *in planta* expression of *liuA* and *liuD* (**Tables S1**) and the iPst19 constrained metabolic flux (**Figure 2**) suggests that leucine, and potentially other BCAAs whose synthesis and degradation are controlled by similar mechanisms (Massey et al., 1976), accumulate to higher levels in bacteria that have been inoculated into MAMP-treated plants compared to those inoculated in mock-treated plants. To test this hypothesis, we supplemented minimal medium with BCAAs, individually or combined, and assessed bacterial growth and gene expression. Supplementation with 10 mM of either leucine, isoleucine, or valine, or the three BCAAs combined, induced the expression of both *liuA* and *liuD* compared to non-supplemented bacteria (**Figure 5A, B**). *liuA* and *liuD* transcriptional responses were specific to BCAAs, as supplementation with 10 mM Gln and Ser had the opposite effect on their expression (**Figure S2**). While leucine had a positive impact on *Pst*DC3000 growth, shortening the doubling time by half, both isoleucine and valine inhibited bacterial growth (**Figure 5C**). When combined, however, the growth-promoting activity of leucine relieved the growth inhibitory effect of isoleucine and valine (**Figure 5D**), suggesting that the three BCAAs serve coordinated functions controlling bacterial metabolism. This is consistent with previous studies showing that changing levels of BCAAs play a regulatory role in bacterial growth and virulence in several gram-positive and gram-negative pathogenic bacteria (Kaiser and Heinrichs, 2018).

**Figure 5:**
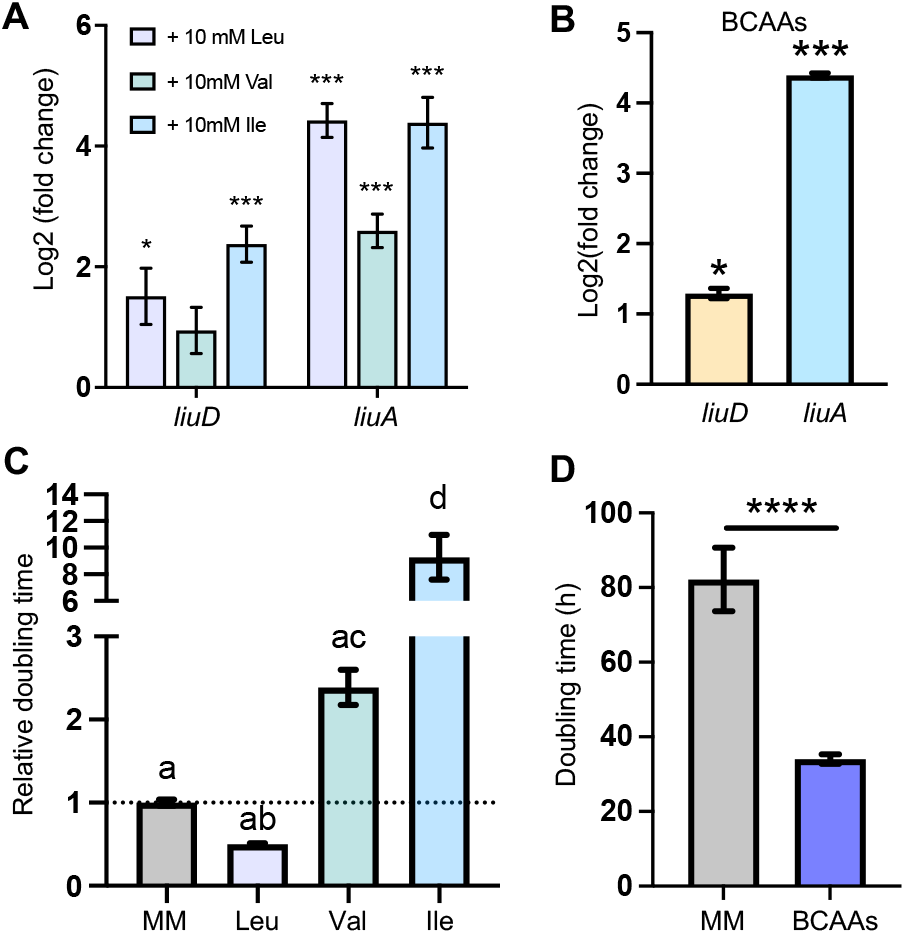
BCAAs levels control both leucine catabolism and PstDC3000 growth *in vitro*. **(A**) Transcript abundance of *liuD* and *liuA* in bacteria exposed to 10 mM Leu, Val, or Ile in HMM normalized to non-supplemented minimal media (N=5). (**B**) Transcript abundance of *liuD* and *liuA* in bacteria exposed to 10 mM each BCAA in HMM normalized to non-supplemented HMM (N=5). (**C**) Bacterial growth in MM9 without supplementation or with 10 mM Leu, Val, or Ile (N= 7-15 cultures). (**D**) Bacterial growth in minimal media without supplementation or with 10 mM of each BCAA.

To test potential connections between BCAA levels and virulence in *Pst*DC3000, we assessed the expression of virulence marker genes in BCAAs supplemented minimal medium. While BCAAs induced the expression of *liuA* and *liuD* (**Figure 5A**), the same amino acids suppressed the expression of the T3SS master regulator gene *hrpL* and the coronatine biosynthesis gene *cfl* when supplemented individually (**Figure 6A**) or combined (**Figure 6B**), suggesting that a BCAAs sensor modulates the expression of BCAA degradation and virulence expression. A similarly low *hrpL* and *cfl* expression was observed 3 h post-inoculation when *Pst*DC3000 was co-infiltrated with BCAAs in naïve plants (**Figure 6C**). In line with mounting evidence suggesting that virulence expression plays an essential role in the early stages of infection (Zhang et al., 2022b), naive plants that are otherwise susceptible to *Pst*DC3000 infections, were able to significantly suppress bacterial growth when co-infiltrated with BCAAs **(Figure 6D**). Importantly, the expression of the salicylic acid and the jasmonic acid plant defense marker genes *PR1* and *VSP2* (Uknes et al., 1992; Berger et al., 2002) was not induced by the infiltration of BCAAs, suggesting that the reduced growth of BCAA-supplemented *Pst*DC3000 is likely due to the direct suppressing effect of BCAA on bacterial virulence expression and not to an induced plant immunity (**Figure S3**).

**Figure 6:**
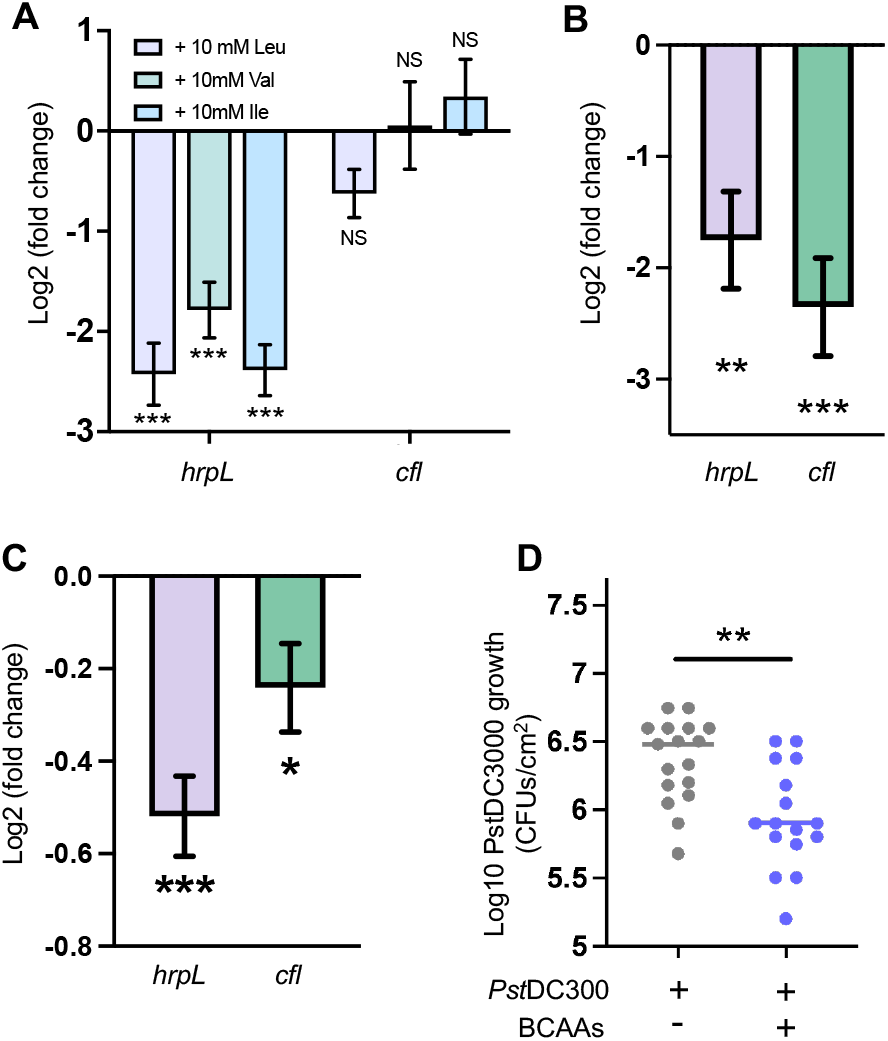
High levels of BCAAs impact the expression of virulence and compromise *Pst*DC3000 colonization of the leaf apoplast. **(A**) Transcript abundance of *hrpL* and *cfl* in bacteria exposed to HMM supplemented with 10 mM of Leu, Val, and Ile in minimal media normalized to non-supplemented HMM (N=5). (**B**) *hrpL* and *cfl* expression in bacteria exposed to HMM supplemented with 10 mM each BCAA normalized to non-supplemented bacteria (N=5). (**C**) *in planta hrpL* and *cfl* expression bacteria co-infiltrated with 10 mM each BCAA and normalized to bacteria with 5 mM MES alone. N = 6. (**D**) PstDC3000 growth 72 HPI in naïve Arabidopsis plants. Bacteria were suspended in 5 mM MES pH 5.7 or 10 mM each BCAA in 5 mM MES pH 5.7 and infiltrated into leaves (N = 17 plants).

### Perturbation of BCAA levels-sensing mechanisms dysregulates virulence gene expression

Changing levels of BCAAs play an important role in coordinating bacterial growth and virulence expression (Pohl et al., 2009; Subashchandrabose et al., 2009). The Leucine-responsive regulatory protein (*Lrp*) is a conserved transcriptional regulator that integrates environmental cues with bacterial metabolism via sensing amino acid levels and coordinating the expression of metabolic and virulence genes in pathogenic bacteria (Kaiser and Heinrichs, 2018). While high levels of BCAAs steer *Lrp* towards transcribing growth-promoting metabolic genes, low levels of BCAAs shift *Lrp*-mediated transcription towards virulence expression, and starvation survival (Tani et al., 2002; Kaiser and Heinrichs, 2018). While the virulence-modulating role of *Lrp*/*AsnC* proteins has been reported for several bacteria species (Baek et al., 2009; Subashchandrabose et al., 2009; Hussa et al., 2015; Casanova-Torres et al., 2017), their functions are largely unknown in *P. syringae*. The *Pst*DC3000 genome encodes nine proteins (**Table S2**) with varying degrees of similarity to the *Escherichia coli Lrp* protein (NP_415409.1). Among these, the protein encoded by the PSPTO_0100 locus is the closest to *E. coli* Lrp with a 61% identity. To test the impact that changes in BCAA levels could have on *Pst*DC3000 metabolism and virulence, we tested bacterial growth and gene expression *in vitro*. In addition, to test the potential regulatory role of *Lrp* in *Pst*DC3000 virulence expression, we overexpressed PSPTO_0100 in sense (*lrp*OX) or antisense (*lrp*AS) orientation (**Figure S4)**. We also made a *Pst*DC3000 control strain that overexpresses the *Arabidopsis uidA* gene (*uidA*OX), a reporter protein, which served as a control to account for the energy expenditure associated with transcribing (*lrpOX* and *lrpAS*) and translating *Lrp* (*lrpOX*) to high levels. Under the virulence-suppressive conditions of a nitrogen-rich medium (Huynh et al., 1989), the *lrp*OX strain expressed significantly more *hrpL* than the control strain (**Figure 7A**). Conversely, the *lrp*AS strain expressed lower than control levels of *hrpL* in HMM (**Figure 7B**), a nutrient-poor medium where *Pst*DC3000 expresses high levels of *hrpL* and other virulence genes (Rahme et al., 1992). The virulence gene *cfl*, essential for the synthesis of the phytotoxin coronatine, showed an expression pattern different from that of *hrpL*. Only in the rich medium was *cfl* expression significantly higher than control levels in *lrp*OX. At the same time, there were no significant differences in *cfl* expression across all strains in HMM (**Figure 7C, D**), suggesting that coronatine biosynthesis and the T3SS are distinctively regulated by *Lrp* activity and hence, BCAA levels.

**Figure 7.**
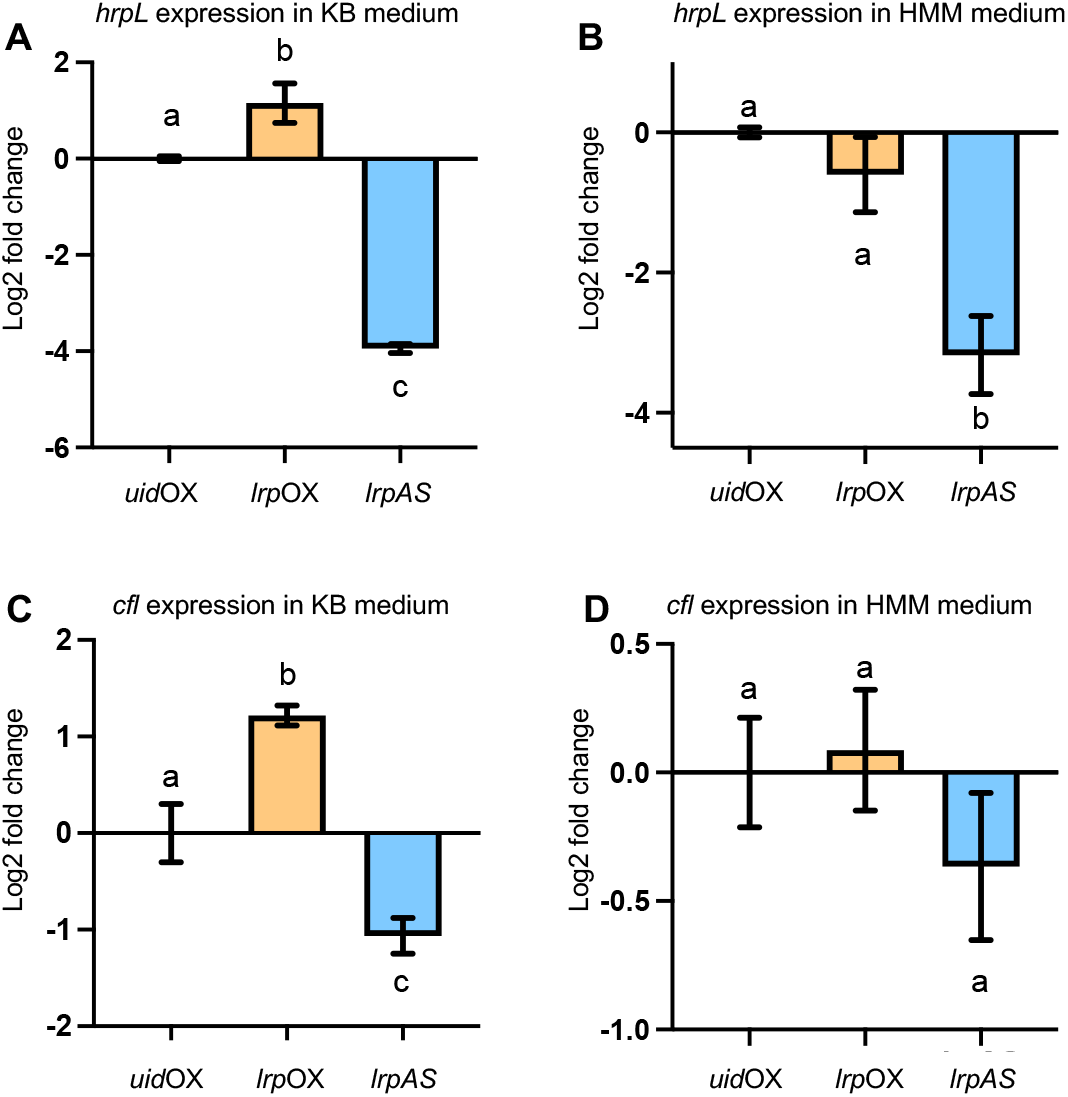
Constitutive Lrp overexpression or down-regulation leads to virulence dysregulation. *hrpL* expression in *lrp*OX and *lrp*AS in KB medium (**A**) or HMM (**B**) at the mid-exponential growth phase normalized to *hrpL* expression in *uid*OX grown in identical conditions. *cfl* expression in *lrp*OX and *lrp*AS in KB medium (**C**) or HMM (**D**) at the mid-exponential growth phase normalized to *cfl* expression in *uid*OX grown in identical conditions. Mean ± SEM (N = 7-13).

## DISCUSSION

### BCAAs modulate *Pst*DC3000 virulence by controlling *Lrp* activity

Branched-chain amino acids serve a central role in integrating extracellular and intracellular cues to maximize bacterial growth and survival (Tani et al., 2002; Kaiser and Heinrichs, 2018). BCAAs exert their regulatory role through the modulation of transcriptional regulators that control the expression of metabolic and virulence genes in pathogenic bacteria (Kaiser and Heinrichs, 2018). Recent studies have provided evidence that BCAAs and the sensor proteins that integrate gene expression also play an important role in the pathogenesis of plant bacterial pathogens. In *Xanthomonas citri* pv. citri, leucine degradation takes place via the acyl-CoA carboxylase complex (ACC) encoded by the *acc* locus. Open reading frames in the *acc* locus show an organization similar to that of the *liu* operon in *P. aeruginosa*. In addition, *AccC* and *AccD* show 53% and 70% identity with *liuD* (PSPTO_2736) and *liuB* (PSPTO_2738), respectively, strongly suggesting that both the *X. citri acc* operon and the *Pst*DC3000 *liu* operon are functionally equivalent and both contribute to leucine catabolism. Mutant strains in *accC* and *accD* showed attenuated growth on citrus plants, suggesting that leucine catabolism in *X. citri* is important for virulence expression *in planta* (Tomassetti et al., 2018). The BCAA sensor protein *Lrp* belongs to a large and conserved family of proteins that bind DNA and regulate gene expression in response to changing levels of intracellular BCAAs (Thaw et al., 2006). The modulation of virulence gene expression exerted by BCAAs on *Lrp* is both ways, positive on certain genes and negative on others (Calvo and Matthews, 1994). In *Erwinia amylovora, Lrp* is necessary to express virulence genes that control motility and synthesis of exopolysaccharides when BCAA levels are low (Schachterle and Sundin, 2019). The evidence contributed by Tomassetti et al. (Tomassetti et al., 2018), Schachterle and Sundin (Schachterle and Sundin, 2019), and the data presented in this study, strongly suggest that leucine degradation, and more broadly, BCAAs degradation, plays a positive role in virulence modulation. The molecular mechanisms by which BCAAs degradation translates into *Lrp*-mediated expression of virulence in these plant pathogens are still unclear. However, our data suggest that a complex balance between *Lrp* protein levels and intracellular levels of BCAAs drives *Lrp* transcriptional activity towards virulence genes expression when *Lrp* is not associated with BCAAs. For instance, the overexpression of *Lrp* induces virulence gene expression in a rich medium where virulence genes are usually suppressed **(Figure 7A**). On the other hand, the *lrp* down-regulation strain (*lrpAS*) shows *hrpL* expression levels below control levels in a minimal medium where *Pst*DC3000 expresses high levels of *hrpL* (**Figure 7B**). In addition, while supplementation with BCAAs induces *liuA* and *liuD* expression, it suppresses the expression of *hrpL* (**Figure 5A, Figure 6A**). Overall, these data suggest that low concentrations of BCAAs allow *Lrp* to express virulence genes. Since high BCAAs levels suppress *Pst*DC3000 virulence gene expression (**Figure 6**) and induce *liuA* and *liuD*, the increased flux in leucine degradation detected in PstDC3000 that have been inoculated in MAMP-treated plants (**Dataset S1**) suggests that lowering BCAAs levels could contribute to re-direct *Lrp* activity towards the expression of virulence genes to counter the overall virulence suppression exerted by MAMP-elicited plants.

### BCAA catabolism in *Pst*DC3000 reactivates virulence expression

In *P. putida*, the *bkd* operon contributes a set of reactions that produce substrates for the *liu* operon (**Figure S1**). Notwithstanding that *Pst*DC3000 lacks a *bkd* operon and possesses a re-arranged *liu* operon, leucine still produces growth when supplied as a carbon source (**Figure 1B; Figure 5**). These data suggest that other enzymes compensate for the loss of the branched-chain ketoacid-dehydrogenases (BCKDH) encoded by the *bdk* operon. In *P. aeruginosa* and *P. putida*, the pyruvate dehydrogenase complex (PDC) and the oxoglutarate dehydrogenase complex (ODHC), share significant homology with the BCKDH enzymes (Burns et al., 1988; Yeaman, 1989; Hester et al., 1995). In *Pst*DC3000, PSPTO_3860, PSPTO_5005, PSPTO_5006, and PSPTO_2201 encode the PDC enzymes, while PSPTO_2199, PSPTO_2200, and PSPTO_2201, encode the ODHC enzymes. Importantly, enzymes in these two complexes show a degree of promiscuity in substrate utilization, suggesting that they could also contribute to BCAAs catabolism in the absence of the *bdk* operon (Heath et al., 2004). Indeed, ODHC enzymes encoded by *Pst*DC3000 showed the highest identity to those encoded by the *bkd* operon in *P. putida* (**Figure S1, Table S2**). It is likely through these enzymes that *Pst*DC3000 still metabolizes leucine as an SCS (**Figure 1B**). Importantly, *Pst*DC3000 synthesizes BCAAs via enzymes encoded by the *ilv* and *leu* operons, both present in its genome (Buell et al., 2003). Notwithstanding the important role of BCAA levels in regulating virulence gene expression, *Pst*DC3000 still responds to changes in the extracellular levels in BCAAs (**Figure 5**) and hence is susceptible to environmental perturbation. In a previous study, we have shown that MAMP-treated plants delay the onset of *Pst*DC3000 virulence via the accumulation of virulence-suppressing amino acids, especially glutamine, serine, and valine (Zhang et al., 2022b), suggesting that the induced expression of *liuA* and *liuD* (**Dataset 1**) would be part of a mechanism that counters the suppression of virulence gene expression. In addition, we showed that 24h post-inoculation in MAMP-treated plants, *Pst*DC3000 expresses *hrpL* to higher levels than in mock-treated plants, suggesting that 24h post-inoculation *Pst*DC3000 has overcome the virulence suppression experienced during the early stages of infection.

Like *Pst*DC3000, *P. aeruginosa* uses BCAA levels to modulate virulence expression. Under low iron conditions that mimic the infection of animal hosts, *P. aeruginosa* induces BCAA catabolism and virulence. This enhanced catabolism leads to a drop in intracellular concentrations of BCAAs that, interestingly, is not compensated by enhanced uptake of BCAA from the environment (Nelson et al., 2019), suggesting that *P. aeruginosa* relies on BCAAs synthesis and degradation to better control virulence and growth. Iron deprivation was a major signature that emerged from the global gene expression analysis of *Pst*DC3000 inoculated in MAMP-treated plants (Nobori et al., 2018). Several of these iron-responsive genes were included in iPst19 (**Dataset S1**), yet through the gene expression-constrained flux, reactions associated with iron metabolism genes did not rank in the top twenty most discordant between mock and flg22 constraints (**Figure 2, Table S1**). Nevertheless, *Pst*DC3000 perceived iron deficiency, together with the increased availability of extracellular amino acids in the LA of MAMP-treated plants, could trigger BCAA degradation and the *Lrp*-mediated re-activation of virulence.

### The power of computational models to uncover the molecular underpinnings of plant-bacteria interactions

The integration of transcriptomic data to constrain iPst19 metabolic flux revealed a metabolic signature that is consistent with previously published studies. While a large proportion of metabolic flux went through glycolysis and the TCA cycle when iPst19 was constrained with bacterial gene expression data from mock-treated plants, leucine catabolism was among the top reactions when iPst19 flux was constrained with bacterial gene expression data from MAMP-treated plants (**Figure 2; Table S1**). This metabolic signature is concordant with the proposed shifts in the composition of LA metabolites described by Yamada et al (Yamada et al., 2016), where MAMP-treated Arabidopsis plants induced the expression of the hexose importers *STP1* and *STP13* that lower hexoses concentrations in the LA. In such a scenario, a lower hexose metabolic activity would be expected in bacteria that were inoculated in MAMP-treated plants compared to mock-treated plants. This prediction was captured by iPst19. The iPst19 constrained flux also predicted the relevance of genes and pathways that were overlooked in previous studies that solely relied on transcriptomics data. *liuA* and *liuD* were among 1294 differentially expressed genes when *Pst*DC3000 was inoculated into MAMP-treated plants (Nobori et al., 2018). Among these 1294 differentially expressed genes, 250 were metabolic genes present in iPst19 (**Dataset S1**). Yet, the constrained metabolic flux analysis indicated that the reactions contributed by most of these 250 genes were not contrasting enough to warrant further study. Importantly, iPst19 rendered a hierarchy of contrasting metabolic reactions that do not correlate with the level of gene expression. In other words, iPst19 predictions were, to some extent, influenced by gene expression. Still, the gene expression integration predicted the relevance of specific genes and pathways that were overlooked in previous studies.

While gene expression analysis reveals important facets of the *Pst*DC3000-Arabidopsis interactions, our findings demonstrate the power of using metabolic computational models to contextualize transcriptomic data. Integration of *in planta Pst*DC3000 transcriptomics has highlighted the role of the BCAAs on virulence gene expression. These predictions were tested and confirmed experimentally, lending support to the strong predictive power of iPst19. In future studies, other transcriptomics datasets could be similarly contextualized using iPst19 to further understand how *Pst*DC3000 metabolism contributes to infecting host plants.

## MATERIALS AND METHODS

### *Pst*DC3000 genome annotations and cross-species comparison

The *Pst*DC3000 genome assembly used was generated by Buell and colleagues (Buell et al., 2003). The draft GENRE was generated using ModelSEED v2.1 (Henry et al., 2010) and the RAST database (Aziz et al., 2008), and further optimized using the COBRApy toolbox (Ebrahim et al., 2013). The *Pst*DC3000 GENRE was refined using cross-species homologous comparisons with two *Pseudomonas aeruginosa* models, iPau1129 and iPae1146. Protein alignments between *Pst*DC3000 features and *P. aeruginosa* features included in their respective GENREs were made with DIAMOND (Buchfink et al., 2014). Comparisons that yielded a significant (e-value < 0.0001) were queried for associated reactions in the *P. aeruginosa* GENREs and subsequently added to the *Pst*DC3000 GENRE with the significantly matching homolog that had functional literature support.

### Ensemble Generation

A full description of the ensemble process and justification was recently published by Medlock and Papin (Medlock and Papin, 2020). iPst19 was generated from the cross-species compared draft reconstruction with integration from single carbon source utilization data (see Biolog growth assays). Each substrate that produced positive growth as defined by significantly different maximum measured OD600nm from the negative control was compiled into a randomly ordered list for use in gap-filling the draft reconstruction. The specific order of metabolites is essential during the gap-filling process, as only the most parsimonious use of the metabolite will result in the addition of a given reaction to the model. If a metabolite can be utilized with the metabolic infrastructure already in place, no new reactions will be added; otherwise, the reactions that add the minimum amount of flux will be added to the model to make use of the metabolite. Only when the draft reconstruction is gap filled and can satisfy the fixed-growth constraint of the biomass function and minimize the fluxes through all other reactions on all *in vitro* growth-producing metabolites (as empirically assessed with Biolog plates), is it then considered a member of the ensemble. The process repeats, starting with the draft reconstruction and a shuffled order of the growth-producing metabolites. As a result of this reshuffling, every member has a slightly different architecture and may produce different biomass fluxes on simulated media. With each round of ensemble gap filling, the 20 most uncertain reactions (where members disagreed on the necessary inclusion of a reaction) introduced by AMMEDEUS we manually curated via literature research. This resulted in the addition of 16 reactions of previously high uncertainty with curated literature support for a particular architecture within the members. The full repository is available at https://github.com/gregmedlock/psy_recon.

### Ensemble Single Gene Deletions

For each member in iPst19, every gene within the member was simulated as a loss of function. For reactions with only one gene association, flux of the reaction became zero. Reactions in which the gene deletion was part of an “and” association, the reaction flux also became zero. For reactions where the deleted gene was in an “or” association, the reaction flux was unaltered. Final readouts of objective function flux were assessed: if the reaction flux was zero or near zero (flux<10e-5), the gene was predicted to be an essential gene.

### Ensemble Multi-Carbon Growth Media Gene Essentiality

Multi-carbon media simulations were made from L-leucine and D-glucose combinations as a percent of the total carbon atoms present in the media. Combinations included 100% glucose, 99% glucose:1% leucine, 90% glucose:10% leucine, 50% glucose:50% leucine, 10% glucose:90% leucine, 1% glucose:99% leucine, and 100% leucine.

### Ensemble Transcriptomic Integration

RNAseq global gene expression profiles of *Pst*DC3000 5h post-inoculation in mock or MAMP-treated plants generated by Nobori and colleagues (Nobori et al., 2018) were integrated into iPst19 members using the GIMME algorithm modified from the Driven package (https://github.com/opencobra/driven). Genes are first stripped from the ensemble, after which they are iteratively added back into the ensemble as a function of expression and significance. The minimum framework needed to satisfy the objective function was assessed for fluxes across all reactions.

### Bacterial Growth Conditions

All bacterial strains were grown in liquid King’s B culture at shaking at 230 RPM and 28°C. Bacteria for growth rate assessment and plant infection experiments were taken from fresh LB agar plates (<1 week old) and grown overnight in the liquid media, followed by sub-culturing 10% until mid-exponential phase.

### Plant growth conditions

*Arabidopsis thaliana* Columbia-0 (Col-0) plants were grown in peat pellets with a 12-hour photoperiod at 23°C and 70% humidity. Plants were watered three times a week with Hoagland’s solution for four weeks and then with water for two more weeks. Six weeks old plants were used for infections and gene expression analysis.

### Bacterial gene expression

For *in vitro* bacterial gene expression, overnight cultures were sub-cultured at 10% in fresh KB media, followed by agitation for two hours at 28°C. Bacteria were pelleted via centrifugation and washed three times in sterile water, after which the bacteria were resuspended to the appropriate OD600 and exposed to experimental conditions. The same method was used to prepare bacteria for *in planta* gene expression assays. The bacteria were resuspended to a final OD600 of 0.2 and pressure-infiltrated into the leaves with a needleless 1 mL syringe. After three hours, infected leaves were harvested and immediately flash frozen in liquid nitrogen; standard RNA isolation from plant tissue or bacteria followed using phenol/chloroform extraction.

### Gene expression analysis

Transcripts of genes were quantified via RT-qPCR. RNA was extracted from leaves or bacterial liquid cultures via phenol/chloroform extraction and treated with DNase 1 (Promega) to remove gDNA contamination. cDNA was synthesized from normalized RNA input using m-MLV reverse transcriptase (Promega). qPCR was performed in duplicate with at least three biological replicates and normalized using the ΔΔCt method.

### Primers, Plasmids, and Bacterial Strains

A complete list of primers (**Table S4**), plasmids, and bacterial strains (**Table S5)** used in this study are available in the supplemental information.

### Plasmid constructs

pBBR5pemIKpKan was purchased from Addgene and prepared via Qiagen miniprep spin kit according to kit specifications. The full-length *lrp* DNA sequence was PCR amplified with Phusion high Fidelity kit (Thermo-Fisher) from *Pst*DC3000 genomic DNA and primers containing attB Gateway recombination site. PCR products were gel purified, and Gateway integrated into pDONR221 (Invitrogen) via BP recombination, followed by LR recombination into pBBR5pemIKpKan to yield pLrpOX and pLrpAS. pENTR-gus (Gateway cloning kit, Invitrogen) was recombined with pBBR5pemIKpKan to yield pUidAOX. *E coli* DH5a competent cells were transformed with the final plasmid constructs selected on gentamicin 25 μg/mL supplemented LB media. Plasmids were transferred to *Pst*DC3000 via triparental mating with helper strain *E. coli* pRK2013.

### Biolog growth assays

Biolog PM1 and PM2 plates were inoculated with 100 μL of IF0 per well, in which *Pst*DC3000 was suspended at 0.07 OD600. Plates were shaken at 7000 rpm and 28°C for 60 hours. OD600 measurements were taken every 12 hours. Breath-Easy sealing membrane was secured to the plate to ensure gas exchange and prevent evaporation.

### *In planta* Bacterial Growth Assays

*Pst*DC3000 was grown as previously described in bacterial growth conditions. Cells were pelleted in microcentrifuge tubes spun at 10,000 RPM. Pellets were washed and resuspended in sterile water to a final inoculation titer of 0.0002 OD600. Six-week-old *Arabidopsis thaliana* Colombia-0 plants were pressure-infiltrated with a needleless 1 mL syringe. Infections proceeded for 72 hours, after which 8 leaf discs were taken with a hole puncher from 4 infected leaves. Discs were homogenized in 400 μL sterile water, with 2 metal beads in 2 mL round bottom tubes in a QIAGEN Tissue Lyser. Ten-fold serial dilutions were plated on LB agar OmniTray plates. CFUs were counted after 16 hours of incubation at 28°C under a dissecting microscope.

## ACKNOWLEDGMENTS

University of Virginia undergraduate students Dianna Quijano and Haziq Mian helped with plant care and bacterial growth, respectively.

---

**Figure S1.**
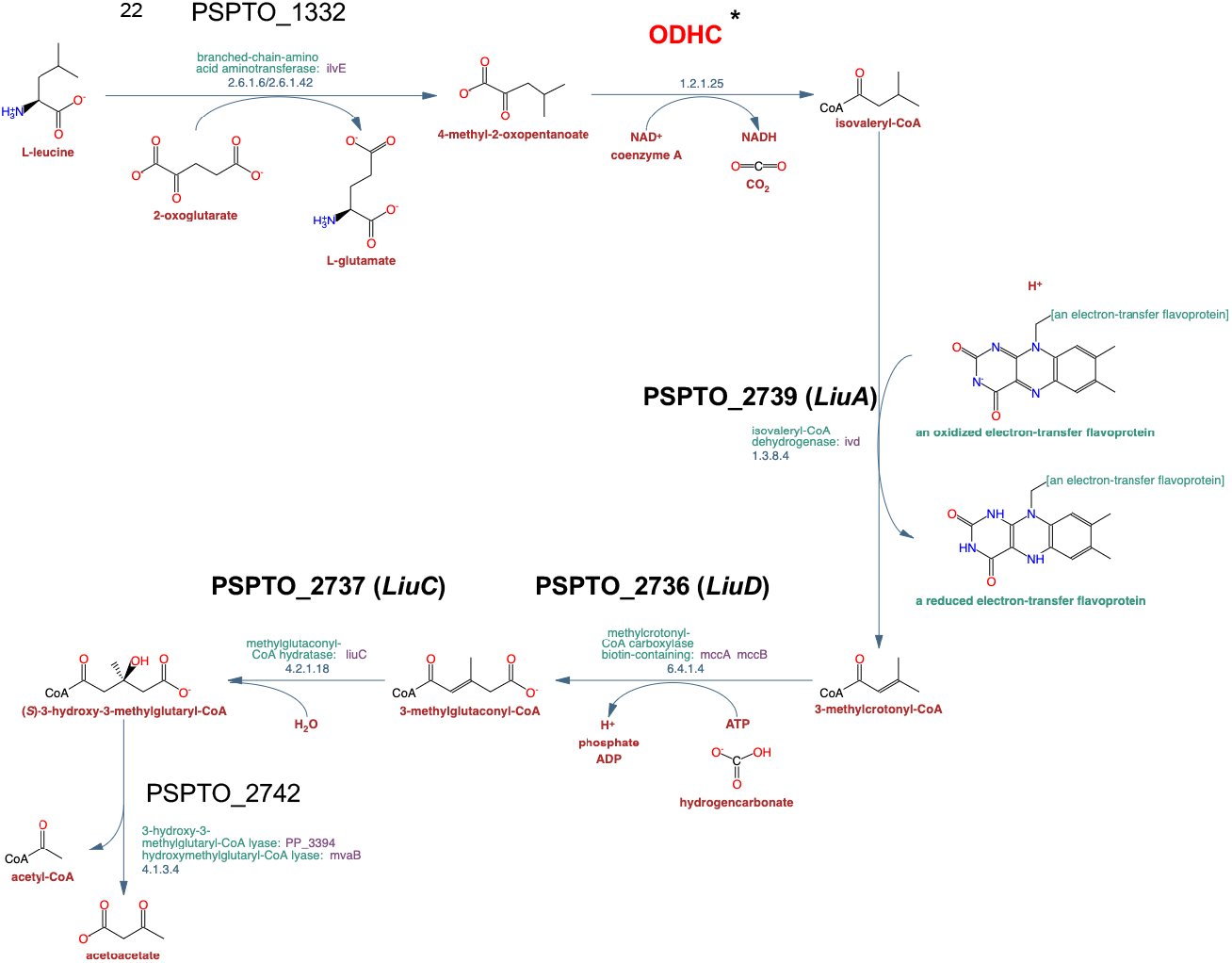
*Pst*DC3000 tentative leucine catabolic pathway. Genes of the *liu* operon are shown in bold. In red is shown the metabolic step tentatively contributed by the *Pst*DC3000 ODHC enzymes. (*) *Pst*DC3000 genes encoding enzymes with significant identity to the BCKDC enzymes encoded by *P. putida bkd* operon: 2-oxoglutarate dehydrogenase-E1 component encoded by PSPTO_2199; 2-oxoglutarate dehydrogenase-E2 component encoded by PSPTO_2200 or PSPTO_5006; 2-oxoglutarate dehydrogenase-E3 component encoded by PSPTO_2201. Modified from MetaCyc (https://metacyc.org/).

**Figure S2:**
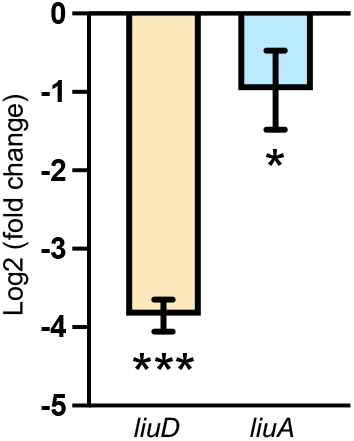
*liuA* and *liuD* expression in response to high levels of glutamine and serine. Transcript abundance of *liuD* and *liuA* in *Pst*DC3000 exposed to 10 mM Gln and Ser in minimal media normalized to non-supplemented minimal media (N = 6).

**Figure S3.**
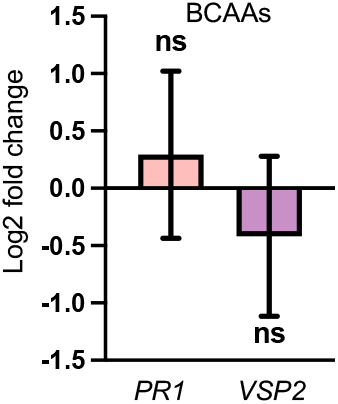
Gene expression of plant defense markers genes *PR1* and *VSP2* 24HPI of leaves with 5 mM MES or 5 mM MES plus 10 mM each BCAA and normalized to 5 mM MES infiltrated leaves (N=3).

**Figure S4:**
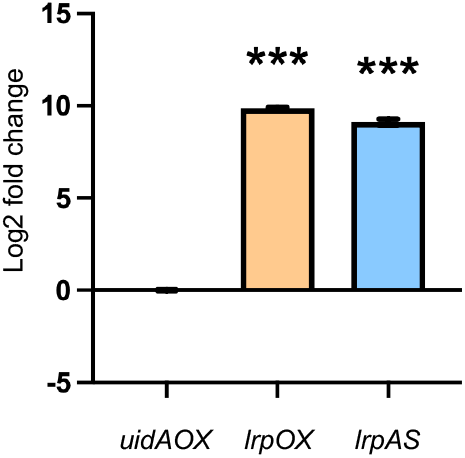
*lrp* expression in *Pst*DC3000 strains harboring either pUidAOX, pLrpOX, or pLrpAS, that overexpress *uidA, and* the *lrp* ORF in the sense orientation (OX) or the antisense (AS) orientation, respectively.

**Table S1:** Metabolic flux analysis constrained with *Pst*DC3000 gene expression *in planta* (Detailed from Figure 2).

| Reaction ID <sup>a</sup> | Reaction Annotation | Pathway | Stoichiometric Equation | GPR <sup>b</sup> | Mock Constraint | Flg22 Constraint | $\Delta^c$ |
| --- | --- | --- | --- | --- | --- | --- | --- |
| EX_cpd00067 | Proton Exchange | - | cpd00067_e <=> | - | -207.9 | -118.86 | -89.04 |
| rxn00512_c | Glycolate:NAD+ oxidoreductase | Glyoxylate and dicarboxylate metabolism | cpd00003_c + cpd00139_c <=> cpd00004_c + cpd00040_c + cpd00067_c | ((PSPTO_2434 or PSPTO_1215) and (PSPTO_1215 or PSPTO_2434)) or (PSPTO_1215 or PSPTO_2434)) | -109.64 | -62.82 | -46.82 |
| EX_cpd00106 | Fumarate Exchange | - | cpd00106_e <=> | - | -47.87 | -12.06 | -35.81 |
| EX_cpd00001 | Water Exchange | - | cpd00001_e <=> | - | 180.53 | 215.86 | -35.33 |
| rxn05319_c | Water Transport | Transport | cpd00001_c --> cpd00001_e | SPONTANEOUS | 180.53 | 215.86 | -35.33 |
| rxn00799_c | (S)-malate hydro-lyase (fumarate-forming) | TCA cycle, Pyruvate Metabolism | cpd00130_c <=> cpd00001_c + cpd00106_c | PSPTO_4339 or PSPTO_1731 or PSPTO_4461 | -33.19 | 0 | -33.19 |
| rxn01388_c | isocitrate hydro-lyase (cis-aconitate-forming) | TCA cycle | cpd00260_c <=> cpd00001_c + cpd00331_c | PSPTO_2016 or PSPTO_3752 | -58.63 | -27.47 | -31.16 |
| rxn05605_c | Maltose permease MAL31 | Transport | cpd00067_e + cpd00130_e <=> cpd00067_c + cpd00130_c | PSPTO_4063 or PSPTO_1682 | 0.6 | 31.05 | -30.45 |
| rxn02866_c | **3-methylbutanoyl-CoA:(acceptor) 2,3-oxidoreductase** | **BCAA metabolism** | **cpd00982_c + cpd01966_c --> cpd00015_c + cpd00067_c + cpd01882_c** | **PSPTO_2739** | **0** | **28.5** | **-28.5** |
| rxn00973_c | citrate hydroxymutase | TCA cycle | cpd00137_c <=> cpd00260_c | PSPTO_3752 or PSPTO_2016 | -27.95 | -2.13 | -25.82 |
| rxn00324_c | glycolate:NADP+ oxidoreductase | Glyoxylate and dicarboxylate metabolism | cpd00006_c + cpd00139_c <=> cpd00005_c + cpd00040_c + cpd00067_c | ((PSPTO_2434 or PSPTO_1215) and (PSPTO_2434 or PSPTO_1215)) | 109.76 | 62.12 | 47.64 |
| rxn00872_c | Butanoyl-CoA:oxygen 2-oxidoreductase | Fatty acid degradation | cpd00650_c + cpd00982_c --> cpd00015_c + cpd00067_c + cpd00120_c | PSPTO_0500 or PSPTO_3706 | 46.61 | 1.84 | 44.77 |
| rxn00781_c | D-glyceraldehyde-3-phosphate:NAD+ oxidoreductase (phosphorylating) | Glycolysis | cpd00003_c + cpd00009_c + cpd00102_c <=> cpd00004_c + cpd00067_c + cpd00203_c | PSPTO_1287 | 46.37 | 1.76 | 44.61 |
| rxn05671_c | D-xylose transport in via proton symport | Transport | cpd00067_e + cpd00154_e <=> cpd00067_c + cpd00154_c | - | 27.5 | -13.44 | 40.94 |
| rxn09650_c | D-xylose transport in via proton symport | Transport | cpd00067_e + cpd00154_e <=> cpd00067_c + cpd00154_c | - | 26.98 | -13.68 | 40.66 |
| rxn00974_c | citrate hydro-lyase (cis-aconitate-forming) | TCA cycle | cpd00137_c <=> cpd00001_c + cpd00331_c | PSPTO_3752 or PSPTO_2016 | 58.63 | 27.47 | 31.16 |
| rxn13659_c | **acyl-CoA dehydrogenase (isovaleryl-CoA)** | **BCAA metabolism** | **cpd00003_c + cpd01882_c <=> cpd00004_c + cpd00067_c + cpd01966_c** | **PSPTO_2739** | **-2.18** | **-30.79** | **28.61** |
| rxn05508_c | Asparagine Transport | Transport | cpd00067_e + cpd00132_e <=> cpd00067_c + cpd00132_c | - | 2.12 | -22.5 | 24.62 |
| EX_cpd00035 | Alanine Exchange | - | cpd00035_e <=> | - | 27.56 | 3 | 24.56 |
| EX_cpd00130 | Malate Exchange | - | cpd00130_e <=> | - | -1.4 | -25.76 | 24.36 |

<sup>a</sup> Reaction IDs from ModelSeed (https://modelseed.org/). <sup>b</sup> Gene-Protein-Reaction. <sup>c</sup> Flux differences between mock and flg22 generated constraints.

**Table S2:** Tentative *Pst*DC3000 genes encoding BCKDC-like enzymes.

| ***P. putida* locus tag**<br>**(query) \*** | ***Pst*DC3000 locus**<br>**tag (hits) \*** | **Identity %** | **E Value** | **Proposed enzyme activity** |
| --- | --- | --- | --- | --- |
| PP_4404 | PSPTO_2201 | 39.65 | 1.70E-102 | ODHC-E3 component |
| PP_4403 | PSPTO_2200 | 32.71 | 6.00E-65 | ODHC-E2 component |
| PP_4402 | PSPTO_2199 | 19.08 | 2.00E-22 | ODHC-E1 component |

(\*) *P. putida* protein sequences were used as query against the *P. syringae* pv. *tomato* DC3000 strain using the DELTA-BLASTp algorithm at NCBI (https://blast.ncbi.nlm.nih.gov/).

**Table S3:** *Lrp* proteins encoded in the *Pst*DC3000 genome

| *Pst*DC3000 locus tag | Protein ID | Protein Name | Query Cover | E value | Identity % * |
| --- | --- | --- | --- | --- | --- |
| PSPTO_0100 | AAO53654.1 | leucine-responsive regulatory protein | 96% | 3.00E-69 | 60.76 |
| PSPTO_3674 | AAO57143.1 | AsnC family transcriptional regulator | 95% | 2.00E-47 | 45 |
| PSPTO_0261 | AAO53807.1 | AsnC family transcriptional regulator | 85% | 4.00E-41 | 42.86 |
| PSPTO_4793 | AAO58223.1 | AsnC family transcriptional regulator | 90% | 1.00E-40 | 44.97 |
| PSPTO_0273 | AAO53819.1 | AsnC family transcriptional regulator | 90% | 3.00E-35 | 36.49 |
| PSPTO_0321 | AAO53866.1 | AsnC family transcriptional regulator | 90% | 4.00E-34 | 34.23 |
| PSPTO_3156 | AAO56642.1 | AsnC family transcriptional regulator | 90% | 5.00E-33 | 35.81 |
| PSPTO_4523 | AAO57971.1 | AsnC family transcriptional regulator | 83% | 3.00E-24 | 27.74 |
| PSPTO_0519 | AAO54062.1 | AsnC family transcriptional regulator | 85% | 2.00E-15 | 27.46 |

(\*) *Lrp* protein sequences from *E. coli* (NP\_415409.1) were used as query against the *P. syringae* pv. *tomato* DC3000 strain using the DELTA-BLASTp algorithm at NCBI (https://blast.ncbi.nlm.nih.gov/).

**Table S4:**
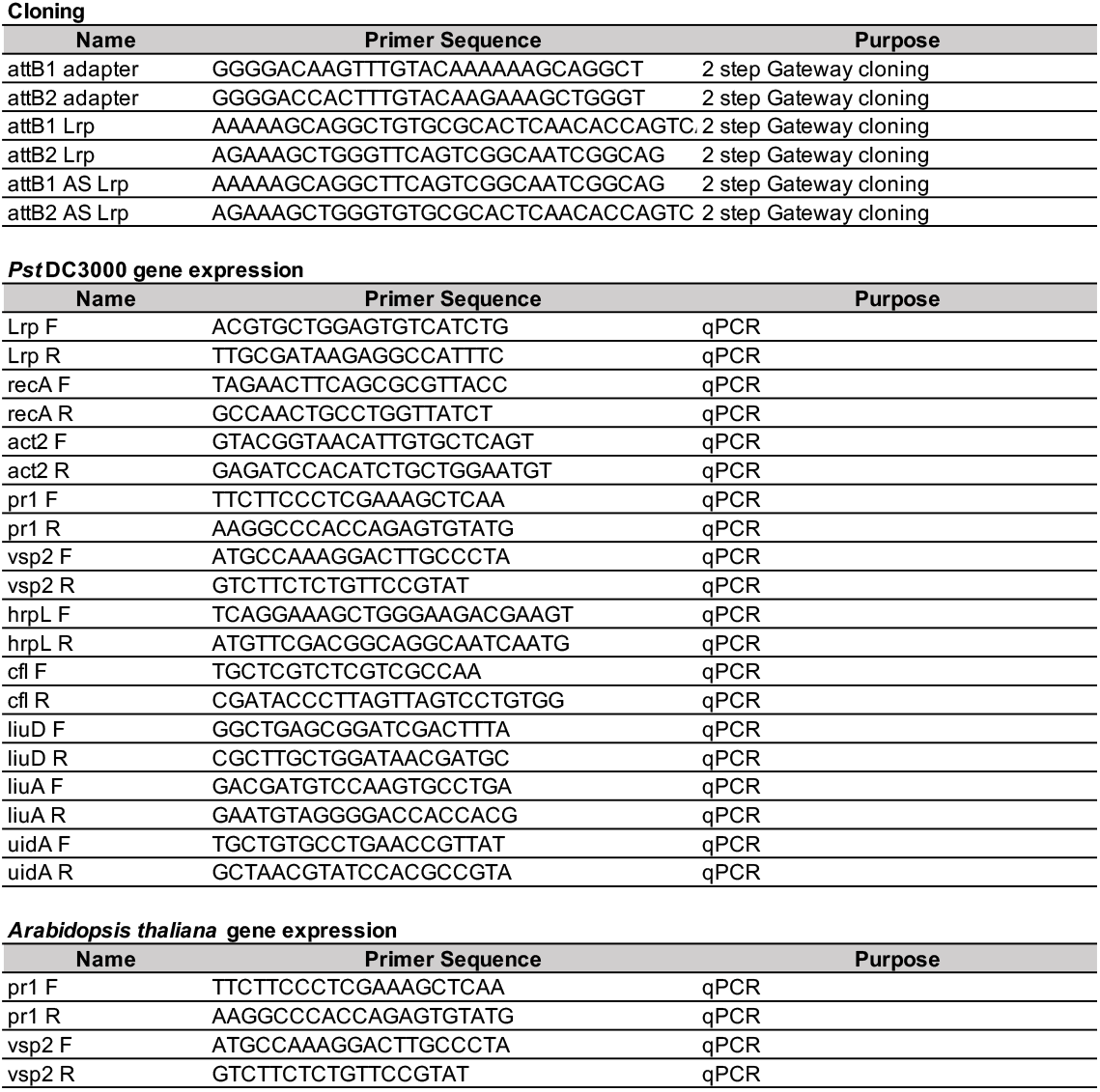
DNA oligonucleotides used in this study

**Table S5:**
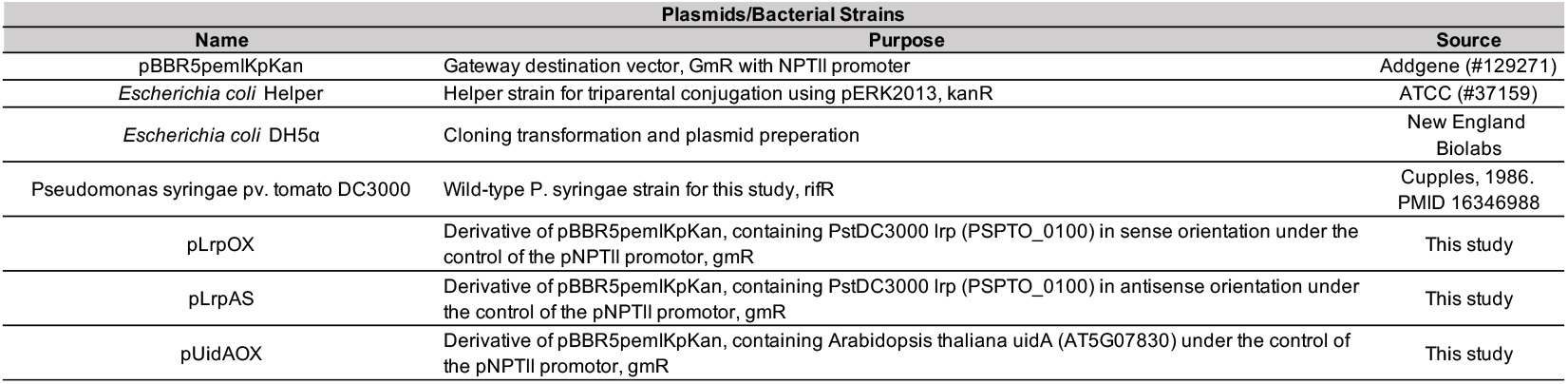
Plasmids and bacterial strains used in this study

| Plasmids/Bacterial Strains |  |  |
| --- | --- | --- |
| Name | Purpose | Source |
| pBBR5pemIKpKan | Gateway destination vector, GmR with NPTII promoter | Addgene (#129271) |
| *Escherichia coli* Helper | Helper strain for triparental conjugation using pERK2013, kanR | ATCC (#37159) |
| *Escherichia coli* DH5α | Cloning transformation and plasmid preparation | New England Biolabs |
| *Pseudomonas syringae* pv. tomato DC3000 | Wild-type *P. syringae* strain for this study, rif<sup>R</sup> | Cupples, 1986.<br>PMID 16346988 |
| pLrpOX | Derivative of pBBR5pemIKpKan, containing PstDC3000 lrp (PSPTO_0100) in sense orientation under the control of the pNPTII promoter, gmR | This study |
| pLrpAS | Derivative of pBBR5pemIKpKan, containing PstDC3000 lrp (PSPTO_0100) in antisense orientation under the control of the pNPTII promoter, gmR | This study |
| pUidAOX | Derivative of pBBR5pemIKpKan, containing Arabidopsis thaliana uidA (AT5G07830) under the control of the pNPTII promoter, gmR | This study |

**Dataset S1:** *Pst* DC30000 gene expression, modified from Nobori et al 2018. Arabidopsis plants were mock-treated or treated with the 22-amino acids synthetic peptide flg22 to elicit plant immunity 24h prior to inoculation with PstDC3000. Bacteria were recovered from inoculated plants 5h after inoculation. Bacterial gene expression was assessed via RNAseq. *liuA* and *liuD* are highlighted in yellow.

| gene_id | log2 fold change<br>flg22_Pst / Mock_Pst | q value |
| --- | --- | --- |
| PSPTO_0062 | 3.415725518 | 8.14E-22 |
| PSPTO_0089 | -1.455019706 | 8.09E-05 |
| PSPTO_0092 | 1.4092938 | 3.58E-15 |
| PSPTO_0103 | 1.237316349 | 1.13E-08 |
| PSPTO_0109 | 1.610088018 | 1.76E-10 |
| PSPTO_0159 | 1.070643278 | 1.03E-11 |
| PSPTO_0218 | 2.022830845 | 2.82E-23 |
| PSPTO_0230 | 1.989391515 | 2.37E-15 |
| PSPTO_0233 | -1.097924212 | 1.12E-05 |
| PSPTO_0259 | 1.343296649 | 0.000138783 |
| PSPTO_0305 | 1.545467794 | 3.58E-20 |
| PSPTO_0308 | 2.723876066 | 5.00E-29 |
| PSPTO_0309 | 1.110449393 | 4.83E-05 |
| PSPTO_0310 | 1.470313706 | 3.70E-05 |
| PSPTO_0311 | 1.112869654 | 0.000114047 |
| PSPTO_0323 | 1.40622995 | 5.24E-20 |
| PSPTO_0386 | -1.120696315 | 8.47E-10 |
| PSPTO_0396 | 2.89309131 | 6.81E-12 |
| PSPTO_0401 | 1.966330018 | 4.46E-10 |
| PSPTO_0455 | 1.957396386 | 2.97E-14 |
| PSPTO_0457 | 2.236705156 | 1.27E-08 |
| PSPTO_0458 | 2.580004255 | 5.14E-22 |
| PSPTO_0460 | 2.349303156 | 8.17E-10 |
| PSPTO_0461 | 2.844178261 | 2.08E-14 |
| PSPTO_0480 | 1.274211573 | 3.53E-15 |
| PSPTO_0495 | -1.237426921 | 4.86E-09 |
| PSPTO_0500 | 1.780579543 | 2.42E-25 |
| PSPTO_0510 | 1.325174806 | 0.001239957 |
| PSPTO_0513 | 1.06555757 | 0.000513285 |
| PSPTO_0518 | 2.194493617 | 3.06E-21 |
| PSPTO_0539 | -1.740594712 | 2.19E-07 |
| PSPTO_0598 | -1.44873455 | 5.66E-10 |
| PSPTO_0622 | -1.041162662 | 9.79E-08 |
| PSPTO_0627 | -1.051813548 | 6.21E-08 |
| PSPTO_0629 | -1.196687896 | 4.77E-15 |
| PSPTO_0631 | -1.504777692 | 3.85E-11 |
| PSPTO_0632 | -1.187877899 | 4.34E-13 |
| PSPTO_0635 | -1.133329053 | 4.01E-07 |
| PSPTO_0641 | -1.056166492 | 2.60E-12 |
| PSPTO_0643 | -1.345004973 | 9.74E-11 |
| PSPTO_0678 | -1.232469943 | 8.23E-09 |
| PSPTO_0692 | -1.066047636 | 5.40E-14 |
| PSPTO_0693 | -1.056843865 | 1.33E-05 |
| PSPTO_0722 | -1.092021429 | 1.00E-06 |
| PSPTO_0726 | 1.823166942 | 2.15E-07 |
| PSPTO_0727 | 2.258863829 | 4.58E-26 |
| PSPTO_0728 | 2.806862516 | 9.24E-43 |
| PSPTO_0744 | 1.80079593 | 4.01E-07 |
| PSPTO_0750 | 2.984836512 | 4.49E-28 |
| PSPTO_0766 | 2.259811382 | 1.07E-21 |
| PSPTO_0774 | 2.408271928 | 6.78E-22 |
| PSPTO_0797 | -1.138088621 | 4.52E-10 |
| PSPTO_0798 | -1.177169177 | 1.23E-08 |
| PSPTO_0800 | 1.02616429 | 2.39E-06 |
| PSPTO_0802 | -1.281151269 | 1.08E-07 |
| PSPTO_0817 | 1.052038758 | 3.92E-08 |
| PSPTO_0873 | -1.531160195 | 2.20E-05 |
| PSPTO_0885 | -2.155417523 | 1.27E-09 |
| PSPTO_0923 | -1.477743621 | 1.87E-06 |
| PSPTO_0957 | 1.099228625 | 1.94E-10 |
| PSPTO_0967 | -1.352928499 | 3.45E-07 |
| PSPTO_1104 | -1.523473474 | 4.66E-15 |
| PSPTO_1105 | -1.292429681 | 4.01E-12 |
| PSPTO_1153 | 2.701656 | 3.09E-11 |
| PSPTO_1182 | 1.59176986 | 7.87E-17 |
| PSPTO_1207 | -1.925310994 | 1.66E-08 |
| PSPTO_1243 | -2.255343662 | 1.53E-20 |
| PSPTO_1292 | 1.881746657 | 1.13E-20 |
| PSPTO_1293 | 1.748373874 | 3.30E-13 |
| PSPTO_1294 | 1.960885291 | 3.96E-16 |
| PSPTO_1295 | 1.215144117 | 2.24E-10 |
| PSPTO_1325 | -1.046872262 | 8.27E-13 |
| PSPTO_1326 | -1.25468978 | 2.06E-09 |
| PSPTO_1329 | -1.392352185 | 1.88E-17 |
| PSPTO_1340 | 1.640209437 | 1.83E-19 |
| PSPTO_1421 | -1.428593493 | 1.89E-07 |
| PSPTO_1428 | -1.367121367 | 1.32E-08 |
| PSPTO_1430 | -1.123231869 | 9.96E-11 |
| PSPTO_1444 | 1.556382476 | 1.90E-23 |
| PSPTO_1450 | -1.290900487 | 1.55E-15 |
| PSPTO_1468 | -1.23987474 | 4.34E-14 |
| PSPTO_1473 | -1.138208649 | 9.05E-18 |
| PSPTO_1501 | -1.048036783 | 2.28E-07 |
| PSPTO_1508 | 1.272689192 | 1.46E-16 |
| PSPTO_1523 | -1.150744637 | 1.10E-06 |
| PSPTO_1538 | -1.881576853 | 2.51E-10 |
| PSPTO_1546 | -1.297912802 | 3.78E-07 |
| PSPTO_1549 | -1.009295935 | 3.13E-14 |
| PSPTO_1552 | -1.10096156 | 1.55E-10 |
| PSPTO_1560 | -2.152800857 | 3.50E-12 |
| PSPTO_1633 | 1.081958614 | 2.54E-09 |
| PSPTO_1661 | -1.334881405 | 7.92E-11 |
| PSPTO_1682 | 1.760415893 | 4.83E-21 |
| PSPTO_1692 | -1.396500381 | 1.57E-07 |
| PSPTO_1700 | -1.003423936 | 2.45E-13 |
| PSPTO_1708 | -1.457180224 | 5.06E-06 |
| PSPTO_1744 | -1.18596107 | 1.61E-06 |
| PSPTO_1748 | -1.824571579 | 5.34E-13 |
| PSPTO_1760 | -1.299417301 | 8.81E-08 |
| PSPTO_1821 | 1.457142609 | 1.77E-10 |
| PSPTO_1825 | 3.64009156 | 7.26E-34 |
| PSPTO_1830 | -1.180715613 | 7.68E-05 |
| PSPTO_1838 | -1.587020925 | 3.97E-08 |
| PSPTO_1859 | 1.538121273 | 2.03E-05 |
| PSPTO_1881 | 1.96946771 | 1.60E-07 |
| PSPTO_1886 | -1.18824405 | 6.39E-09 |
| PSPTO_1890 | 3.219548745 | 4.28E-12 |
| PSPTO_1907 | 1.690114586 | 4.98E-06 |
| PSPTO_1930 | 1.170509847 | 1.10E-06 |
| PSPTO_2016 | 1.324809557 | 4.51E-10 |
| PSPTO_2026 | 2.23324503 | 9.81E-13 |
| PSPTO_2046 | -1.341201946 | 1.69E-06 |
| PSPTO_2055 | -1.10430127 | 5.42E-09 |
| PSPTO_2136 | -2.546452162 | 3.47E-18 |
| PSPTO_2141 | -1.514194171 | 5.55E-09 |
| PSPTO_2173 | 1.384740308 | 1.06E-21 |
| PSPTO_2174 | 1.417797077 | 3.02E-16 |
| PSPTO_2177 | 1.630906256 | 9.34E-15 |
| PSPTO_2178 | 1.759678311 | 8.22E-12 |
| PSPTO_2179 | 1.516553296 | 4.28E-09 |
| PSPTO_2206 | -1.019556734 | 0.00089864 |
| PSPTO_2338 | 2.80919509 | 2.61E-21 |
| PSPTO_2341 | 2.007763533 | 2.61E-09 |
| PSPTO_2367 | 1.010970474 | 4.34E-06 |
| PSPTO_2371 | 1.147775913 | 0.002279982 |
| PSPTO_2396 | 3.316806521 | 6.68E-17 |
| PSPTO_2397 | 2.740984574 | 7.15E-19 |
| PSPTO_2401 | 3.486447676 | 4.71E-17 |
| PSPTO_2510 | 1.038839979 | 1.25E-05 |
| PSPTO_2636 | 2.730554475 | 2.54E-31 |
| PSPTO_2638 | 2.398657239 | 1.26E-27 |
| PSPTO_2639 | 2.112570104 | 4.42E-15 |
| PSPTO_2655 | 2.353167766 | 1.88E-07 |
| PSPTO_2656 | 1.054719597 | 0.000983925 |
| PSPTO_2668 | 1.326075099 | 2.02E-06 |
| PSPTO_2673 | 1.19698883 | 1.13E-05 |
| PSPTO_2701 | 2.215108441 | 4.44E-14 |
| PSPTO_2702 | 2.702451969 | 7.48E-20 |
| PSPTO_2703 | 2.314053669 | 1.16E-14 |
| PSPTO_2705 | 2.862564933 | 1.25E-14 |
| PSPTO_2707 | 3.167745289 | 2.49E-25 |
| PSPTO_2723 | 2.040235158 | 7.62E-08 |
| PSPTO_2736 | 3.607111983 | 3.20E-20 |
| PSPTO_2739 | 3.473796666 | 1.10E-27 |
| PSPTO_2760 | 1.403011965 | 6.16E-09 |
| PSPTO_3002 | 1.809773893 | 8.22E-09 |
| PSPTO_3062 | 1.100338675 | 8.74E-06 |
| PSPTO_3125 | 1.883779385 | 4.56E-18 |
| PSPTO_3149 | -1.434749517 | 4.06E-15 |
| PSPTO_3155 | 3.187327695 | 2.69E-25 |
| PSPTO_3163 | 1.727432179 | 1.63E-06 |
| PSPTO_3164 | 2.573206523 | 3.89E-14 |
| PSPTO_3262 | 2.770040464 | 1.30E-16 |
| PSPTO_3344 | -1.724533072 | 1.01E-09 |
| PSPTO_3360 | -1.261339506 | 3.49E-17 |
| PSPTO_3364 | 2.676408123 | 1.28E-28 |
| PSPTO_3455 | 3.405377805 | 3.81E-31 |
| PSPTO_3489 | 3.976092858 | 2.77E-22 |
| PSPTO_3494 | 3.373472376 | 3.26E-20 |
| PSPTO_3516 | 1.582946198 | 1.16E-13 |
| PSPTO_3517 | 1.713955855 | 3.07E-21 |
| PSPTO_3550 | 1.932875457 | 2.46E-26 |
| PSPTO_3551 | 1.78622407 | 2.16E-25 |
| PSPTO_3553 | 1.298124207 | 4.45E-18 |
| PSPTO_3559 | 1.553621653 | 3.28E-14 |
| PSPTO_3593 | -1.506987975 | 3.97E-10 |
| PSPTO_3660 | 2.104174329 | 1.23E-20 |
| PSPTO_3661 | 1.880330794 | 3.50E-21 |
| PSPTO_3662 | 1.456293096 | 1.02E-10 |
| PSPTO_3675 | 2.74211639 | 7.90E-11 |
| PSPTO_3705 | 1.072756062 | 7.24E-11 |
| PSPTO_3806 | 2.050075064 | 8.27E-11 |
| PSPTO_3811 | -1.023914482 | 2.77E-10 |
| PSPTO_3814 | -1.402635182 | 5.71E-06 |
| PSPTO_3816 | -1.246749997 | 3.58E-07 |
| PSPTO_3817 | -1.332751597 | 5.52E-07 |
| PSPTO_3829 | -1.961859773 | 9.97E-12 |
| PSPTO_3835 | -1.59766798 | 1.90E-08 |
| PSPTO_3950 | -1.209159512 | 1.21E-07 |
| PSPTO_4098 | 1.586463016 | 4.36E-11 |
| PSPTO_4108 | 1.251117899 | 6.03E-22 |
| PSPTO_4168 | 1.237213408 | 3.19E-07 |
| PSPTO_4170 | 1.965901193 | 1.76E-11 |
| PSPTO_4179 | 4.065072208 | 8.37E-32 |
| PSPTO_4307 | 1.816416371 | 2.77E-10 |
| PSPTO_4312 | -1.388001695 | 4.55E-15 |
| PSPTO_4397 | -1.591151783 | 7.57E-07 |
| PSPTO_4399 | -1.543735176 | 5.46E-15 |
| PSPTO_4425 | -1.426081244 | 1.03E-07 |
| PSPTO_4461 | -2.351939767 | 6.92E-14 |
| PSPTO_4494 | -1.591779773 | 8.87E-08 |
| PSPTO_4503 | -1.642186647 | 4.11E-08 |
| PSPTO_4547 | 1.662228702 | 1.41E-17 |
| PSPTO_4549 | 2.865702838 | 1.44E-25 |
| PSPTO_4650 | 1.287498618 | 1.52E-08 |
| PSPTO_4686 | -2.693307419 | 1.69E-40 |
| PSPTO_4687 | -2.775878676 | 2.07E-35 |
| PSPTO_4799 | -1.434429663 | 8.03E-08 |
| PSPTO_4800 | -1.103389253 | 8.19E-11 |
| PSPTO_4814 | -1.559024062 | 1.21E-08 |
| PSPTO_4825 | -1.42370877 | 7.62E-13 |
| PSPTO_4828 | -1.269749907 | 7.81E-06 |
| PSPTO_4866 | -1.571559178 | 2.62E-23 |
| PSPTO_4867 | -1.290927122 | 5.99E-09 |
| PSPTO_4874 | -1.212641698 | 1.84E-07 |
| PSPTO_4876 | -1.513236219 | 3.98E-09 |
| PSPTO_4877 | -1.362678564 | 4.66E-07 |
| PSPTO_4918 | 1.144819386 | 0.000616064 |
| PSPTO_4919 | 2.068970435 | 1.32E-11 |
| PSPTO_4991 | -2.002752685 | 1.59E-10 |
| PSPTO_5005 | -1.119918151 | 5.17E-18 |
| PSPTO_5060 | 3.173584138 | 4.69E-30 |
| PSPTO_5064 | 1.928083775 | 1.15E-15 |
| PSPTO_5110 | -1.106458236 | 5.34E-05 |
| PSPTO_5134 | -1.183510222 | 1.52E-14 |
| PSPTO_5138 | -1.307851075 | 1.53E-15 |
| PSPTO_5171 | 2.228269925 | 6.06E-14 |
| PSPTO_5188 | 1.606760701 | 4.12E-08 |
| PSPTO_5227 | -1.434997576 | 6.48E-06 |
| PSPTO_5231 | 2.394324182 | 1.31E-12 |
| PSPTO_5255 | 1.962905487 | 1.10E-16 |
| PSPTO_5270 | 2.784087388 | 1.97E-26 |
| PSPTO_5274 | 2.090461658 | 2.81E-13 |
| PSPTO_5278 | 1.416457657 | 5.17E-11 |
| PSPTO_5279 | 2.852235507 | 2.26E-27 |
| PSPTO_5289 | -1.977501909 | 1.65E-10 |
| PSPTO_5308 | 1.175377278 | 1.97E-10 |
| PSPTO_5314 | 1.562134165 | 2.45E-12 |
| PSPTO_5316 | 2.591033248 | 7.65E-25 |
| PSPTO_5319 | 2.276813515 | 2.79E-14 |
| PSPTO_5335 | -1.507147074 | 4.03E-10 |
| PSPTO_5338 | -1.80112545 | 1.41E-08 |
| PSPTO_5356 | 2.316294658 | 5.75E-23 |
| PSPTO_5357 | -1.463132325 | 1.46E-08 |
| PSPTO_5358 | 1.319743529 | 1.04E-11 |
| PSPTO_5360 | -1.093868987 | 0.000267142 |
| PSPTO_5378 | 1.296179943 | 3.74E-05 |
| PSPTO_5393 | 1.294277682 | 1.71E-07 |
| PSPTO_5406 | 1.464508001 | 5.24E-12 |
| PSPTO_5461 | 2.671721676 | 1.78E-19 |
| PSPTO_5501 | -1.178278284 | 3.91E-05 |
| PSPTO_5510 | 2.497097877 | 1.17E-14 |
| PSPTO_5511 | 2.738005067 | 2.63E-23 |
| PSPTO_5530 | 1.066838825 | 2.73E-06 |
| PSPTO_5595 | 1.456942463 | 2.91E-21 |
| PSPTO_5598 | -1.457055038 | 6.17E-10 |
| PSPTO_5600 | -1.27385437 | 3.18E-21 |
| PSPTO_5603 | -1.18717908 | 1.32E-09 |
| PSPTO_5604 | -1.180141811 | 9.46E-06 |
| PSPTO_5614 | -1.761104743 | 2.80E-07 |

